# Limited window for donation of convalescent plasma with high live-virus neutralizing antibodies for COVID-19 immunotherapy

**DOI:** 10.1101/2020.08.21.261909

**Authors:** Abhinay Gontu, Sreenidhi Srinivasan, Eric Salazar, Meera Surendran Nair, Ruth H. Nissly, Denver Greenawalt, Ian M. Bird, Catherine Herzog, Matthew J. Ferrari, Indira Poojary, Robab Katani, Scott E. Lindner, Allen M. Minns, Randall Rossi, Paul A. Christensen, Brian Castillo, Jian Chen, Todd N. Eagar, Xin Yi, Picheng Zhao, Christopher Leveque, Randall J. Olsen, David W. Bernard, Jimmy Gollihar, Suresh V. Kuchipudi, James M. Musser, Vivek Kapur

## Abstract

The optimal timeframe for donating convalescent plasma to be used for COVID-19 immunotherapy is unknown. To address this important knowledge deficit, we determined *in vitro* live-virus neutralizing capacity and persistence of IgM and IgG antibody responses against the receptor-binding domain and S1 ectodomain of the SARS-CoV-2 spike glycoprotein in 540 convalescent plasma samples obtained from 175 COVID-19 plasma donors for up to 142 days post-symptom onset. Robust IgM, IgG, and viral neutralization responses to SARS-CoV-2 persist, in the aggregate, for at least 100 days post-symptom onset. However, a notable acceleration in decline in virus neutralization titers ≥160, a value suitable for convalescent plasma therapy, was observed starting 60 days after first symptom onset. Together, these findings better define the optimal window for donating convalescent plasma useful for immunotherapy of COVID-19 patients and reveal important predictors of an ideal plasma donor, including age and COVID-19 disease severity score.

**One Sentence Summary:** Evaluation of SARS-CoV-2 anti-spike protein IgM, IgG, and live-virus neutralizing titer profiles reveals that the optimal window for donating convalescent plasma for use in immunotherapy is within the first 60 days of symptom onset.

## MAIN TEXT

The kinetics and longevity of the antibody response to severe acute respiratory syndrome coronavirus 2 (SARS-CoV-2) are poorly understood. This knowledge is essential for determining if individuals have been infected, elucidating host and virus factors that influence the magnitude and persistence of serological responses, assessing whether an individual is sufficiently protected from re-infection, and evaluating the effectiveness of vaccination strategies to contain the pandemic. Additionally, understanding antibody kinetics and persistence is essential to determine correlates of live-virus neutralization (VN) titers required for qualifying donors of convalescent plasma for use in immunotherapy.

Antibodies directed to the SARS-CoV-2 surface spike glycoprotein (S) ectodomain (S/ECD) and receptor-binding domain (S/RBD) neutralize SARS-CoV-2 *in vitro*, and their titers can serve as effective surrogates for virus neutralization (VN)^1–3^. These titers have also been used to identify suitable convalescent plasma donors for COVID-19 immunotherapy^1,3^. However, there is considerable uncertainty about the robustness and persistence of the serological responses to SARS-CoV-2. Some reports suggest variable duration and resilience of serum IgG or IgM antibodies to S or other viral proteins^2–4^, whereas others report that serological and neutralizing responses begin to wane and approach undetectable levels within weeks after infection^3–6^.

To better understand the kinetics of the serological response to SARS-CoV-2, we determined the temporal profiles of IgM, IgG, and VN responses in a cohort of 175 convalescent plasma donors, including 105 who had donated multiple times. Plasma samples (*n=*540) were collected up to 142 days after the onset of the donors’ first symptoms [days post-symptom onset (DPO); Tables 1,S1]. We used a Fab fragment-based assay to assess total antibody titers against S/ECD and S/RBD, an isotype-specific assay to measure anti-S/RBD IgM and IgG titers, and a live-virus assay to determine SARS-CoV-2 VN titers^1^.

**Table 1:** Demographics and characteristics of the plasma donor cohort.

| Patient characteristics | Samples<br>n (%) | Individuals<br>n (%) |
| --- | --- | --- |
| **Sex** |  |  |
| Female | 213 (39.4) | 88 (50.3) |
| Male | 327 (60.6) | 87 (49.7) |
| **Age** |  |  |
| 20-30 | 95 (17.6) | 26 (14.9) |
| 31-40 | 117 (21.7) | 39 (22.3) |
| 41-50 | 166 (30.7) | 51 (29.1) |
| 51-60 | 117 (21.7) | 40 (22.9) |
| > 60 | 45 (8.3) | 19 (10.9) |
| Average (95% CI) | 43.8 (42.7 - 44.9) | 44.9 (43.0 - 46.8) |
| Median (IQR) | 44 (33 - 53) | 46 (36 - 54) |
| Range | 20 - 78 | 20 - 78 |
| **Severity** |  |  |
| 1 | 244 (45.2) | 76 (43.4) |
| 2 | 182 (33.7) | 63 (36.0) |
| 3 | 23 (4.3) | 10 (5.7) |
| 4 | 44 (8.1) | 15 (8.6) |
| 5 | 47 (8.7) | 11 (6.3) |
| Median (IQR) | 2 (1 - 2) | 2 (1 - 2) |
| Range | 1 - 5 | 1 - 5 |
| **Dyspnea** |  |  |
| No | 250 (46.3) | 79 (45.1) |
| Yes | 290 (53.7) | 96 (54.9) |
| **DPO** |  |  |
| < 31 | 39 (7.2) | 35 (20.0) |
| 31-60 | 181 (33.5) | 89 (50.9) |
| 61-90 | 173 (32.0) | 44 (25.1) |
| 91-120 | 122 (22.6) | 7 (4.0) |
| > 120 | 25 (4.6) | - |
| Average (95% CI) | 70.8 (68.4 - 73.3) | 49.5 (46.5 - 52.5) |
| Median (IQR) | 68 (48 - 93) | 46 (32 - 63) |
| Range | 17 - 142 | 17 - 108 |
| **Hospitalization** |  |  |
| No | 428 (79.3) | 141 (80.6) |
| Yes | 112 (20.7) | 34 (19.4) |
| **Total** | 540 | 175 |

We discovered robust IgM, IgG, and VN responses in the majority of individuals, with moderate to strong correlation regardless of assay type (Figure 1a,b). Only 4 of 175 [2.3%; 95% confidence interval (CI): 0.9–5.7%] individuals had undetectable levels of IgG, IgM, or total antibody to S/RBD or S/ECD at initial sampling, whereas a significantly higher fraction (29 of 114; 25.4%; 95% CI: 18.3-34.1%) had undetectable VN titers (z-score=6; *P*<0.01). Thus, ~75% of RT-PCR-confirmed symptomatic individuals were serologically positive for anti-spike protein antibody, and their convalescent plasma had demonstrable ability to neutralize SARS-CoV-2 in VN assays.

**Figure 1.**
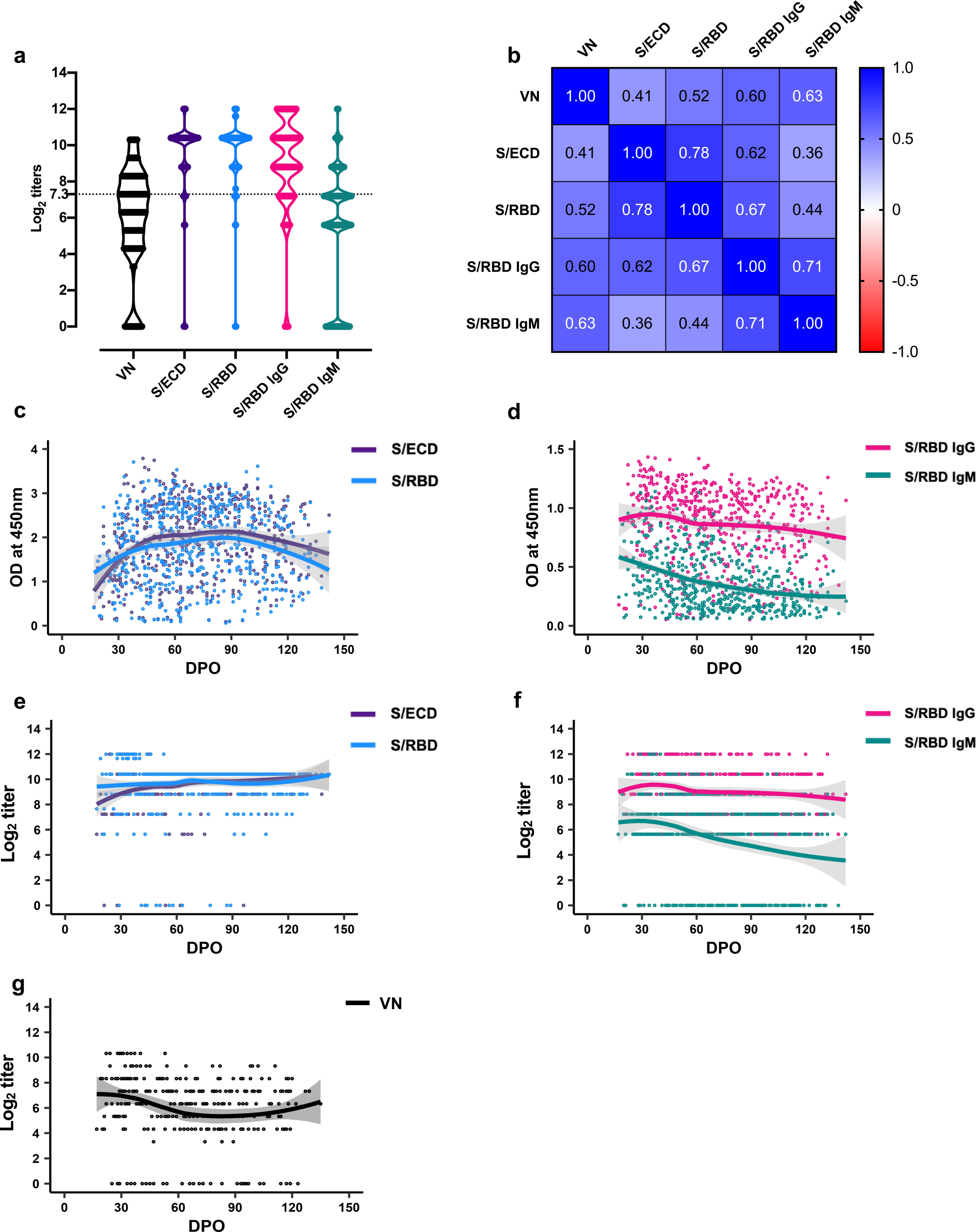
Distribution, correlation, and trajectories of antibody titers against SARS-CoV-2. (a) Violin plots showing distribution of virus neutralization titers (*n*=305); total antibody (*n*=538), and specific isotype antibody IgG and IgM (*n*=540) titers to SARS-CoV-2 spike-ectodomain (S/ECD) and spike-receptor binding domain (S/RBD) in convalescent plasma samples (Log_2_ transformed values). The means of the distribution among the titers were significantly different, except between S/ECD and S/RBD [One-way ANOVA, Tukey’s multiple comparison (mixed-effects model), *P*<0.05]. The dashed line at Log_2_ titer represents VN titer of 1:160. (b) Pairwise comparison of the assays show a moderate to strong correlation between the total and isotype specific IgG and IgM antibody estimates with virus neutralization assays. (c) & (d) Optical density (OD) (at 450nm) for the indirect ELISAs indicating total or isotype specific IgG and IgM antibody levels; (e) & (f) Titers of the total or isotype specific IgG and IgM antibodies. The IgG and IgM titers appear to peak around 30 days post onset (DPO) of symptoms. High IgG titers persist until 140 DPO, while IgM titers trend lower but persist until 140 DPO. (g) Neutralizing antibody titers persist until 140 DPO. A locally estimated scatterplot smoothing (LOESS) regression curve is fitted to the data.

We next determined the patterns of distribution of IgM and IgG background-corrected optical density (OD) values and titers over time (Figure 1c-f). Titers peaked at approximately 30 DPO and persisted through 140 DPO (Figure 1c-f), with the IgG titer consistently higher than the IgM titer. The titer ratios began to diverge after 60 DPO (Figure 1d,f), but remained strongly correlated over the first 140 DPO (Pearson’s *r*=0.71; 95% CI: 0.67–0.75). The observed persistence of IgG responses in many convalescent individuals through 140 DPO is encouraging from the perspective of antibody durability to SARS-CoV-2. The data are consistent with the expected serological responses to rapidly replicating RNA viruses, including SARS-Cov-1^7^. In contrast, the persistence of IgM well beyond the acute phase was unexpected and differs from reports suggesting a rapid decline in IgM by 4-6 weeks^7,8^.

To further study the trajectory of antibody persistence, we performed survival analyses on IgM and IgG titers on all 540 samples obtained from 175 individual donors (Figure 2). Consistent with the temporal distribution of titers, survival analyses showed that the proportion of S/RBD IgG Fc seropositive convalescent individuals remained high through 140 DPO (Figure 2a,b).

**Figure 2.**
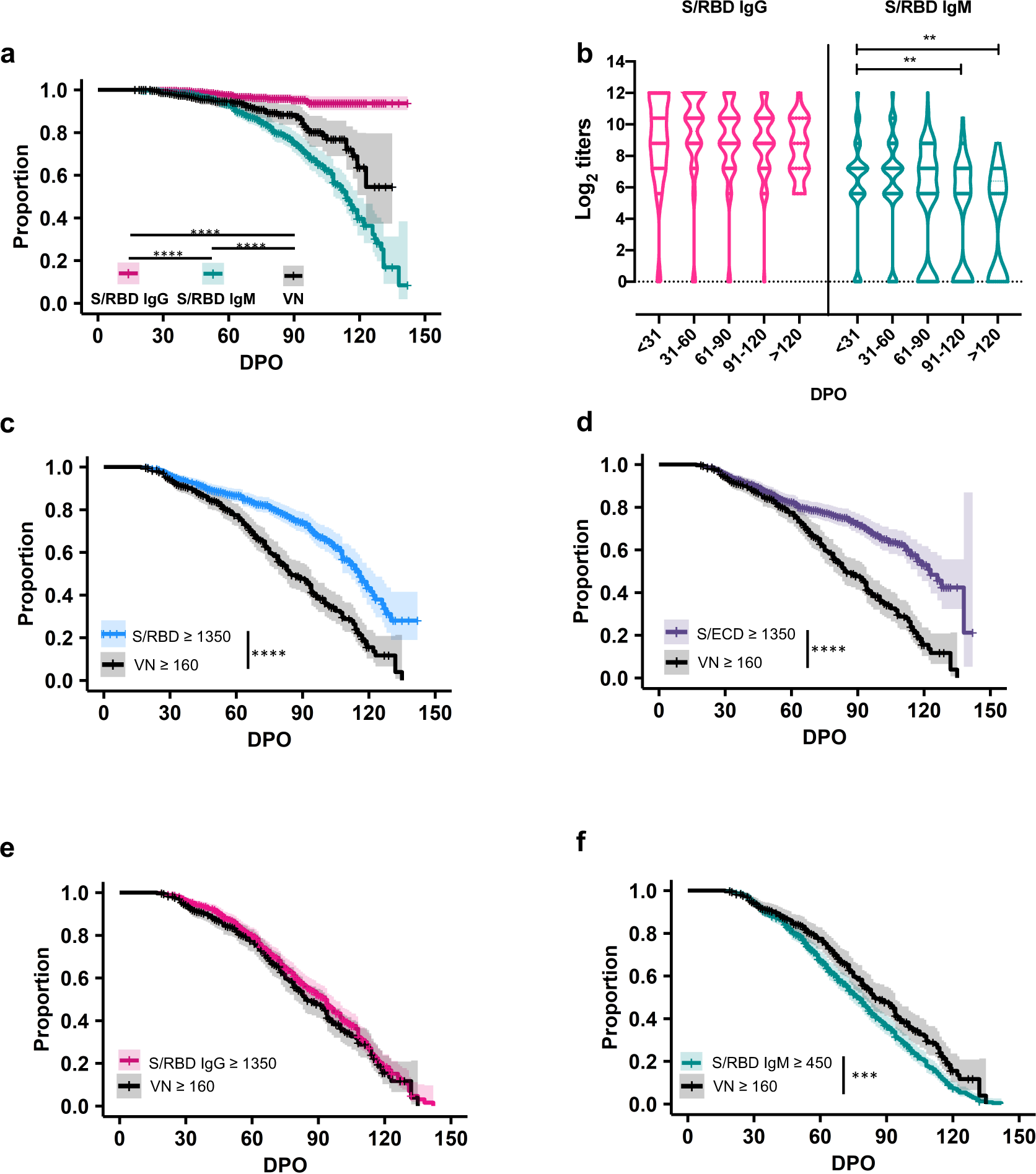
Survival analysis of IgG and IgM antibody titers to SARS-CoV-2 spike-receptor binding domain (S/RBD) in 540 samples and virus neutralizing antibody (VN) titers in 305 samples collected from convalescent individuals (*n*=175) during the first 140 days post onset of symptoms (DPO). (a) Proportion of S/RBD IgG seropositive convalescent individuals remains high through 140 DPO, while IgM seropositivity remains high through the first 60 DPO and then steadily declines over the next 60 days (Log rank test; \*\*\*\**P*<0.0001). The proportion of individuals with VN responses also begins to decline 60 DPO, with ~50% of individuals remaining seropositive with VN test through 140 DPO (Log rank test; \*\*\**P*<0.001). (b) Violin plots showing a significant decline in VN and IgM titers with time (Ordinary one-way ANOVA, Tukey’s multiple comparison test; \**P*<0.05; \*\**P*<0.01); the IgG titers remain stable until after 120 DPO. Comparison of proportion of individuals seropositive with S/RBD, S/ECD, and S/RBD IgG titers ≥1350 as well as with S/RBD IgM titer ≥450 to the proportion of individuals possessing VN titers ≥160 through 140 DPO are depicted in c, d, e, and f respectively (\*\*\**P*<0.001; \*\*\*\**P*<0.0001).

It is clear that antibodies directed against SARS-CoV-2 S/ECD and S/RBD neutralize the virus *in vitro*. Consistent with this, several vaccines targeting the S glycoprotein have shown promise in animal infection models and human clinical trials^9–13^. We and others have recently reported that anti-S/RBD and S/ECD IgG titers are excellent surrogates for *in vitro* VN and help identify plasma donors for therapeutic uses^1,14^. Specifically, we have shown that anti-S/RBD or anti-S/ECD antibody titers of ≥1350 are strong proxies for a VN titer ≥160, the FDA-recommended value for use in COVID-19 convalescent plasma therapy^1^, and transfusion of anti-S/RBD IgG ≥1350 titer plasma within 72 hours (h) of hospitalization significantly improves survival and health outcomes^15,16^.

Our large and well-characterized convalescent plasma library with longitudinally donated samples enabled detailed assessment of VN response persistence. We found that the proportion of individuals with a VN titer ≥160 remained above 80% through the first 60 DPO but declined sharply to less than 20% between DPO 61 and 120 (Figure 2c). These results suggest that the time period in which donated convalescent plasma is likely to have a high VN titer and optimal therapeutic potential is within the first 60 DPO. This has important implications for convalescent plasma donation and passive immunotherapy programs, some of which have already transfused more than 60,000 individuals in the United States as of August 13, 2020 (https://www.uscovidplasma.org).

Facile methods to identify suitable convalescent plasma donors are needed as the gold standard live-virus VN assays used herein are labor intensive, cumbersome, take several days to perform, and require specialized expertise and access to a high containment (Biosafety Level 3) laboratory and regulatory clearances. ELISAs are easier to implement than VN assays, especially in resource-limited countries and environments. We previously reported that an S/RBD ≥1350 titer may serve as a good marker for identifying plasma donors with VN ≥160^1^ (Supplementary Table S2). Here we confirm a high positive likelihood ratio (LR+; 13.43) for a VN ≥160 when S/RBD titers are ≥1350 early (1-30 DPO) post onset of symptoms (Supplementary Table S2). However, extended longitudinal analyses through 140 DPO show that S/ECD and S/RBD ≥1350 persist longer than VN ≥160, with significantly different survival curves (*P<*0.001) for 1-140 DPO and overall LRs+ of 1.34 for S/ECD and 1.61 for S/RBD (Figure 2c,d; Supplementary Table S2). Thus, an S/RBD ≥1350 titer is a promising marker for identifying suitable plasma donors early, but not late, after first symptom onset. In contrast, S/RBD IgG ≥1350 appears to be a reliable predictor of VN ≥160, and S/RBD IgG ≥1350 survival is statistically indistinguishable from that of VN ≥160 (Figure 2e), with an overall LR+ of 3.18 and a negative likelihood ratio (LR-) of 0.26 (Supplementary Table S2).

We next investigated the survival and predictive values of S/RBD IgM ≥450 as compared to VN ≥160 (Figure 2f, Supplementary Table S2). An S/RBD IgM titer ≥450 was selected because the magnitude of IgM response was approximately three-fold lower than that of IgG (Figure 1f). The results showed that S/RBD IgM ≥450 had a similar survival profile to VN ≥160 but waned significantly faster (*P <*0.01; Figure 2f). While S/RBD IgM ≥450 had an overall LR+ of 3.72, it also had a LR-of 0.69, which would likely result in an unacceptable number of suitable donors with VN ≥160 being excluded. Together, these results indicate that S/RBD IgG ≥1350, but not IgM ≥450 or S/RBD or S/ECD total antibody ≥1350, serves as a good marker to identify suitable plasma donors for COVID-19 immunotherapy.

To determine the kinetics and persistence of IgM, IgG, and VN responses, we next performed longitudinal analyses of the initial and final observed titers in 105 subjects with multiple plasma donations [median 4 donations, interquartile range (IQR): 2-6; median interval between initial and final donation of 42 days (range 6-101; IQR: 26-68), Extended Data Figure 1]. The data confirm the robustness of IgG and IgM levels through the 140 DPO observation period. All individuals with a detectable starting titer remained, on average, between one or two dilutions above or below the initial titer (Extended Data Figure 1). Of particular note, only 5 of 60 individuals (8.3%, 95% CI: 2.8-18.4%) with an initial VN titer of ≥ 5.3 (1:40) showed a subsequent increase in titer, emphasizing the importance of recruiting and screening convalescent plasma donors quickly, as VN titers are unlikely to rise from initial levels.

We next assessed whether particular donor characteristics predicted a more robust serological and neutralization response. The results show that individuals 30 years of age or younger had significantly lower VN, IgG and IgM antibody titers than those in the older age groups (Figure 3a). Individuals between ^20–30^ years of age also had significantly faster decline in IgG (*P <0.05*) and IgM (*P <0.05*) than did those > 60 years of age (Figure 3b-d, Extended Figure 2a). Consistent with recent evidence that disease severity correlates with the magnitude and duration of serological response^1,17,18^, we found that individuals with disease severity scores of 4 or 5 on a 5-point disease severity scale had significantly higher IgM and IgG antibody titers than those with lower severity scores (Figure 3e). In addition, survival analyses of IgG and IgM antibody titers revealed that individuals with mild/moderate symptoms scores of 1, 2, or 3 had significantly different survival curves for IgM (*P<*0.001) and VN (*P<*0.05) than did those with higher disease severity scores (3f-h, Extended Figure 2b). Notably, all individuals with high severity scores had detectable IgM at their last measurement point, as did all individuals who were >60 years of age. This may be indicative of potential confounding or interaction between age and disease severity affecting the magnitude and persistence of serological response. The rate of loss of IgM seropositivity to S/RBD was significantly higher for the youngest (20-30 years) compared to the oldest (>60 years) age groups (log-rank test, *P*<0.01), and this effect remained significant when individuals with high severity scores were excluded. Age and severity score were only weakly correlated (Spearman rank correlation=0.08; *P*=.07), but formal analysis of confounding or interactions between age and severity was precluded due to data frailty and requires further study. Regardless, these findings suggest that convalescent individuals <30 years of age and those with lower disease severity scores are less likely to represent suitable donors of convalescent plasma for immunotherapy for COVID-19 patients than individuals in >30 age group with a history of more severe disease. Finally, the results show that individuals with dyspnea had significantly higher VN, IgG and IgM and antibody titers than those who did not (Figure 3i), and IgM seropositivity declined significantly faster in individuals with dyspnea (log-rank test, *P*<0.0001) (Figure 3j-l).

**Figure 3.**
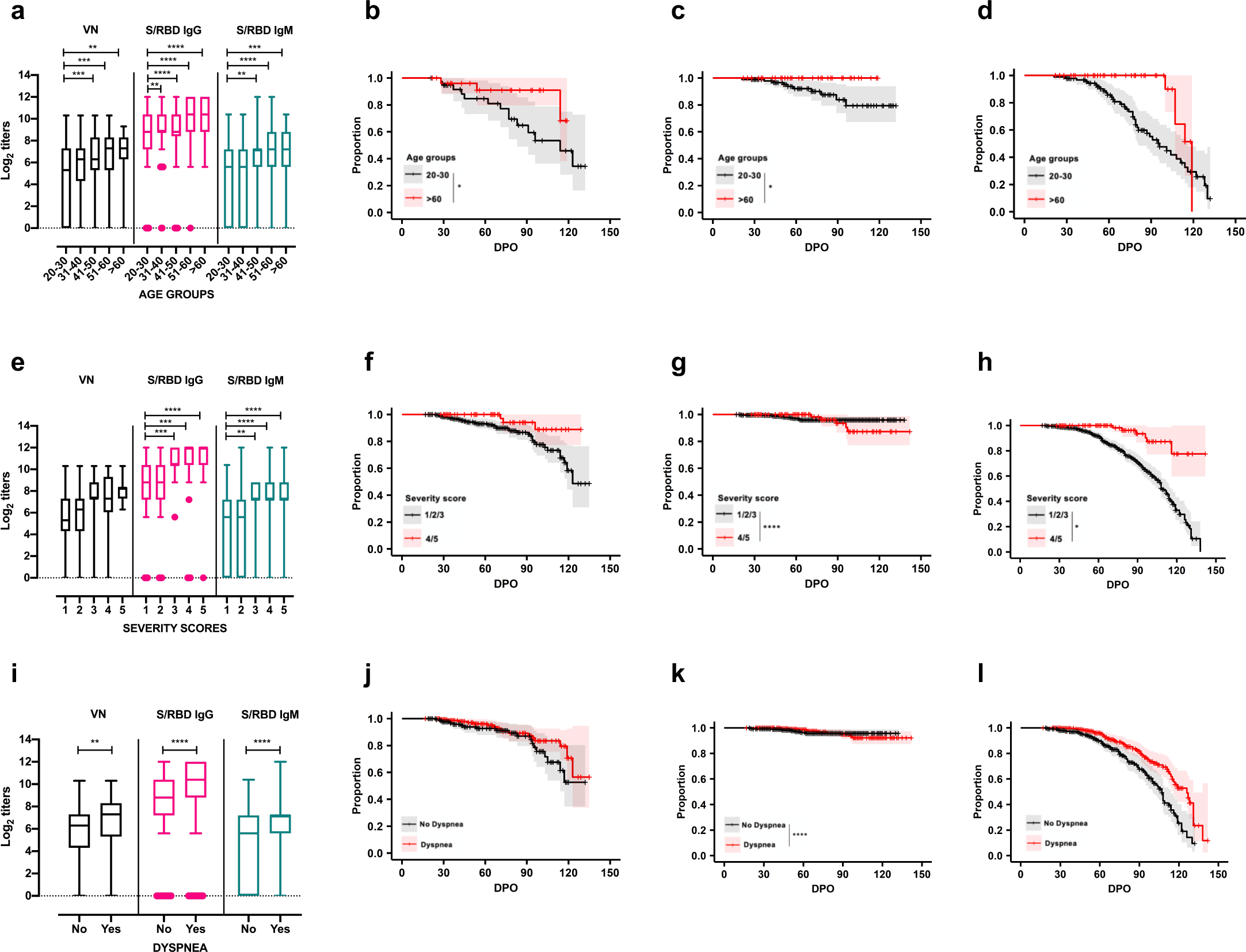
Distribution of antibody titers against SARS-CoV-2 based on age, severity scores, and presence of dyspnea. These data represent samples collected from convalescent individuals (*n*=175) during the first 140 days post symptom onset (DPO). (a) Individuals <31 years of age have significantly lower IgG, IgM, and viral neutralizing antibody (VN) titers than those >40 years of age in this cohort (Ordinary one-way ANOVA, Tukey’s multiple comparison test; \*\**P*<0.01; \*\*\**P*<0.001; \*\*\*\**P*<0.0001). Survival analysis of (b) IgG, (c) IgM, and (d) VN antibody titers during the first 140 DPO in convalescent individuals within the age groups of 20-30 (*n*=95 samples) and >60 (*n*=45 samples) (Log-rank test, \**P*<0.05 for IgG and IgM, *P*>0.05 for VN antibodies). (e) Individuals with a severity score of 1 have significantly lower IgM and IgG titers than those above a score of 3 (Ordinary one-way ANOVA, Tukey’s multiple comparison test; \*\**P*<0.01; \*\*\**P*<0.001; \*\*\*\**P*<0.0001). Survival analysis of (f) IgG, (g) IgM, and (h) VN antibody titers in relation to severity scores grouped as mild (1/2/3) and severe (4/5) in convalescent individuals during the first 140 DPO (Log-rank test, *P*>0.05 for IgG, \*\*\*\**P*<0.0001 for IgM, \**P*<0.05 for VN antibodies). (i) Individuals with dyspnea had significantly higher VN, IgM, and IgG titers (Ordinary one-way ANOVA, Tukey’s multiple comparison test; \*\**P*<0.01; \*\*\*\**P*<0.0001). Survival analysis of (j) IgG, (k) IgM, and (l) VN antibody titers in relation to occurrence of dyspnea in convalescent individuals during the first 140 DPO (Log-rank test, *P*>0.05 for IgG, \*\*\*\**P*<0.0001 for IgM, *P*>0.05 for VN).

In conclusion, these data refine our understanding of the kinetics, magnitude, and durability of human serologic responses to SARS-CoV-2 spike protein, the primary vaccine candidate being studied worldwide. This integrative analysis of serological and VN profiles identifies an optimal donation window of up to 60 DPO for high-titer anti-spike protein convalescent plasma as immunotherapy for COVID-19 patients. Our analysis found that additional characteristics of an ideal potential donor include a recovered patient >30 years old with a high COVID-19 disease severity score. In the aggregate, these data permit a more focused strategy for identifying suitable donors for COVID-19 convalescent plasma and passive immunotherapy programs.

## Online Methods

### Data Availability

All data generated or analyzed during this study are included in this published article (and its supplementary information files) or will be made available by the authors on reasonable request.

#### Cohort and sample description

Plasma samples (n=540) from 175 COVID-19 convalescent patients collected at Houston Methodist Hospital in Houston, Texas were included in the study. Patients were confirmed to be positive for SARS-CoV-2 by RT-PCR. The severity of infection in these patients was scored on a scale of 1-5, (median 2, IQR: 1-2). Clinical improvement relative to DPO 0 was defined as a 1 point improvement in ordinal scale [1, discharged (alive); 2, not hospitalized, experiencing dyspnea not requiring supplemental oxygen but requiring ongoing medical care (for COVID-19 or otherwise); 3, hospitalized, requiring low-flow supplemental oxygen; 4, hospitalized, on non-invasive ventilation or high-flow oxygen devices; 5, hospitalized and on invasive mechanical ventilation or extracorporeal membrane oxygenation (ECMO)].

Per FDA guidelines (https://www.fda.gov/vaccines-blood-biologics/investigational-new-drug-ind-or-device-exemption-ide-process-cber/recommendations-investigational-covid-19-convalescent-plasma#Patient%20Eligibility), all subjects were asymptomatic for at least 14 days at the time of plasma collection. Of the 175 subjects, 105 eligible individuals underwent plasmapheresis and donated plasma at least twice (range 2-12 times). All donors were confirmed negative for SARS-CoV-2 by RT-PCR and provided written consent before plasmapheresis. The study cohort consisted of 88 females (50.3%) and 87 males (49.7%), ranging in age between ^20–78^ years (median 46, IQR: 36-54). Samples were collected from 17-142 DPO (median 68 days, IQR: 48-93). Plasma from donors was collected with the transfusion apheresis system (Trima Accel^®^ Terumo BCT) and standard blood banking protocols were followed. An aliquot of collected plasma was tested for antibodies by ELISA and/or VN assays. Cohort characteristics are described in Table 1 and Supplementary Table 1.

#### Study approvals

Informed consent was obtained from either the patient or an authorized representative of the patient when applicable for collection of plasma samples. All procedures were approved by the Institutional Review Board of Houston Methodist Hospital (IRB# PRO00025121). Serological analyses were performed at the Pennsylvania State University under BSL-2 (ELISA assays) and BSL-3 (VNs) conditions, following the Pennsylvania State University Institutional Biosafety Committee (IBC) approved protocols.

#### Quantitative estimation of antibodies against SARS-CoV-2

SARS-CoV-2 antibodies in plasma samples were detected and quantified against purified recombinant SARS-CoV-2 spike ectodomain (S/ECD) or receptor-binding domain (S/RBD) proteins using in-house indirect Fab antibody-based or isotype-specific (IgM and IgG) ELISA assays. The protocols were performed as previously described^1,19^ and deposited in protocols.io (dx.doi.org/10.17504/protocols.io.bivgke3w). Two isotypes of CR3022, a human monoclonal antibody reactive to spike regions of SARS-CoV-1 and SARS-CoV-2, were used as positive controls in the assays (IgG1: Ab01680-10.0; IgM: Ab01680-15.0, Absolute Antibody, USA). The cutoff for the assays was determined as an optical density (absorbance at 450 nm) higher than three or six standard deviations above the mean of the tested pre-COVID-19 serum samples (*n*=100). Sample titers were estimated as reciprocals of the highest dilution resulting in an OD greater than the cutoff. The class specificity of the IgM ELISA was tested by treating the plasma samples (*n*=10) with 1,4-Dithiothreitol (DTT, 10708984001, Millipore Sigma, USA) as previously described^20^. Briefly, samples were allowed to react with 0.005 M DTT in PBS at 36±2°C for 30 min and then tested with isotype-specific ELISAs for titer estimation (Extended data Figure 3).

#### Virus neutralization assay

The VN titers of the plasma samples were quantified on a cell-based assay using SARS-CoV-2 strain USA-WA1/2020 (NR-52281-BEI Resources, USA) based on procedures described previously^1,21^. Briefly, Vero E6 cells (CRL-1586, ATCC, USA) were grown as monolayers in 96-well microtiter plates. Heat-inactivated plasma samples were diluted two-fold in triplicate and incubated with 100 tissue culture infective dose 50 (TCID_50_) of the virus at 5% CO_2_ at 36±2°C for 60 min. This plasma-virus mixture was added to cell monolayers and incubated further for 72 h at 5% CO_2_ at 36±2°C. Plates were treated with crystal violet formaldehyde stain for 1 h and visually inspected for cytopathic effect (CPE) or protection. The reciprocal of the highest dilution of the plasma where at least two of the three wells were protected (no CPE) was determined as the VN titer of the sample.

#### Statistical analyses

Tests for normality were performed using the Kolmogorov-Smirnov test and a *P* value of <0.05 was considered statistically significant. Data dispersion was indexed by standard errors of mean or quartile and IQR. The agreement between the various assays was determined using Pearson correlation coefficient with log2-transformed titers. The non-parametric regression method LOESS was used for scatterplot smoothing to visualize antibody trajectories. The geom_smooth (method=”loess”) function in R was used with default span of 0.75. The proportion of the sample population remaining seropositive over the 100-day period was determined using a log-rank test and Kaplan-Meier survival curves were plotted with “survival” and “survminer” packages in R Studio^22–24^. Statistical differences in antibody titers and survival curves of patient characteristics—including severity score, age, and presence of dyspnea—were analyzed using one-way ANOVAs (Tukey’s multiple comparison tests) and a log-rank test, respectively. Individual level interval-censored data were used to fit semi-parametric accelerated failure time models using the icenReg R package. DTComPair R package (https://cran.r-project.org/web/packages/DTComPair/DTComPair.pdf) was used to compare the sensitivity, specificity, and positive and negative predictive values for detection of S/RBD, S/ECD, and S/RBD IgG titers ≥1350, as well as S/RBD IgM titer ≥450 using VN titer ≥160 as the gold standard. Positive and negative predictive values were compared with the generalized score statistics, whereas the sensitivity and specificity were compared using an exact binomial test. All analyses were completed using R (versions 3.6.1 or 3.6.3) within R Studio (version 1.2.5019) or Graphpad PRISM 8 (version 8.4.3).

## Acknowledgments

We are deeply indebted to all of our volunteer plasma donors for their time, their generous gift, and their solidarity. We thank Katharine G. Dlouhy, Curt Hampton, and their team of coordinators and recruiters for outstanding efforts; and Monisha Dey, Cheryl Chavez-East, John Rogers, Dr. Ahmed Shehabeldin, Dr. David Joseph, Guy Williams, Karen Thomas, and Curt Hampton who were instrumental in efficiently managing the donor center; Drs. Jessica Thomas and Zejuan Li, Erika Walker, the very talented and dedicated molecular technologists, and the many labor pool volunteers in the Molecular Diagnostics Laboratory for their dedication to patient care; the many donor center and blood bank phlebotomists and technologists for their dedication to donor and blood safety; Drs. Heather McConnell and Sasha M. Pejerrey for outstanding editorial assistance; Brandi Robinson, Harrold Cano, and Cory Romero for technical assistance; Claude Moussa, Heather Patton, and the many members of the laboratory information technology team for rapidly implementing the necessary electronic workflows; Pamela McShane, Dilzi Mody, and the many members of the biorepository team for their meticulous management of patient samples; and Christina Talley, Dr. Susan Miller, and Mary Clancy for consistent, thorough, and outstanding advice. We express our gratitude to Manuel Hinojosa and Mark Vassallo for their extensive efforts to rapidly procure resources, and Dr. Roberta Schwartz for her efforts in implementing screening of asymptomatic individuals. We are indebted to Drs. Marc Boom and Dirk Sostman for their support, and to many very generous Houston citizens and businesses for their tremendous philanthropic support of this ongoing project, including but not limited to anonymous, Ann and John Bookout III, Carolyn and John Bookout, Ting Tsung and Wei Fong Chao Foundation, Ann and Leslie Doggett, Freeport LNG, the Hearst Foundations, Jerold B. Katz Foundation, C. James and Carole Walter Looke, Diane and David Modesett, the Sherman Foundation, Paula and Joseph C. “Rusty” Walter III, and Aramco Americas. Dr. Jason S. McLellan (University of Texas at Austin) graciously provided the mAb CR3022 and the spike protein expression vectors, and we thank the members of the Center for Systems and Synthetic Biology at the University of Texas at Austin for technical assistance. We thank Zivko Nikolov, Susan Woodard, and Michael Johanson at the National Center for Therapeutics Manufacturing at Texas A&M University for production of antigen. We thank Terumo BCT for continuously and rapidly supplying blood collection devices and supplies, and Victoria Cavener and Team COVID-19 serology at Penn State for their timely and generous technical assistance and logistical support. This study was supported by the Fondren Foundation, Houston Methodist Research Institute (to JMM). This research has been funded in whole or part with federal funds under a contract from the National Institute of Allergy and Infectious Diseases, National Institutes of Health, Contract Number 75N93019C00050 (to JL and GCI). A portion of this work was funded through Cooperative Agreement W911NF-12-1-0390 by the Army Research Office (to JDG). We gratefully acknowledge the unwavering support and timely seed funding from the Huck Institutes of the Life Sciences for the studies at Penn State together with the Huck Distinguished Chair in Global Health award (to VK).

## Author Contributions

Project concept (AG, SS, ES, SVK, JMM, VK); acquired data (AG, SS, ES, MSN, RHN. DG, IMB, IP, RK, SEL, AMM, RR, PAC, BC, JC, TNE, XY, PZ, CL, RJO, DWB, JG); analyzed data (AG, SS, MSN, CH, MJF, SVK, JMM, VK); wrote manuscript (VK, JMM, AG, SS, SVK); prepared figures (AG, SS, MSN, CH, VK). All authors reviewed the manuscript and gave final approval for publication.

---

**Extended Data Figure 1.**
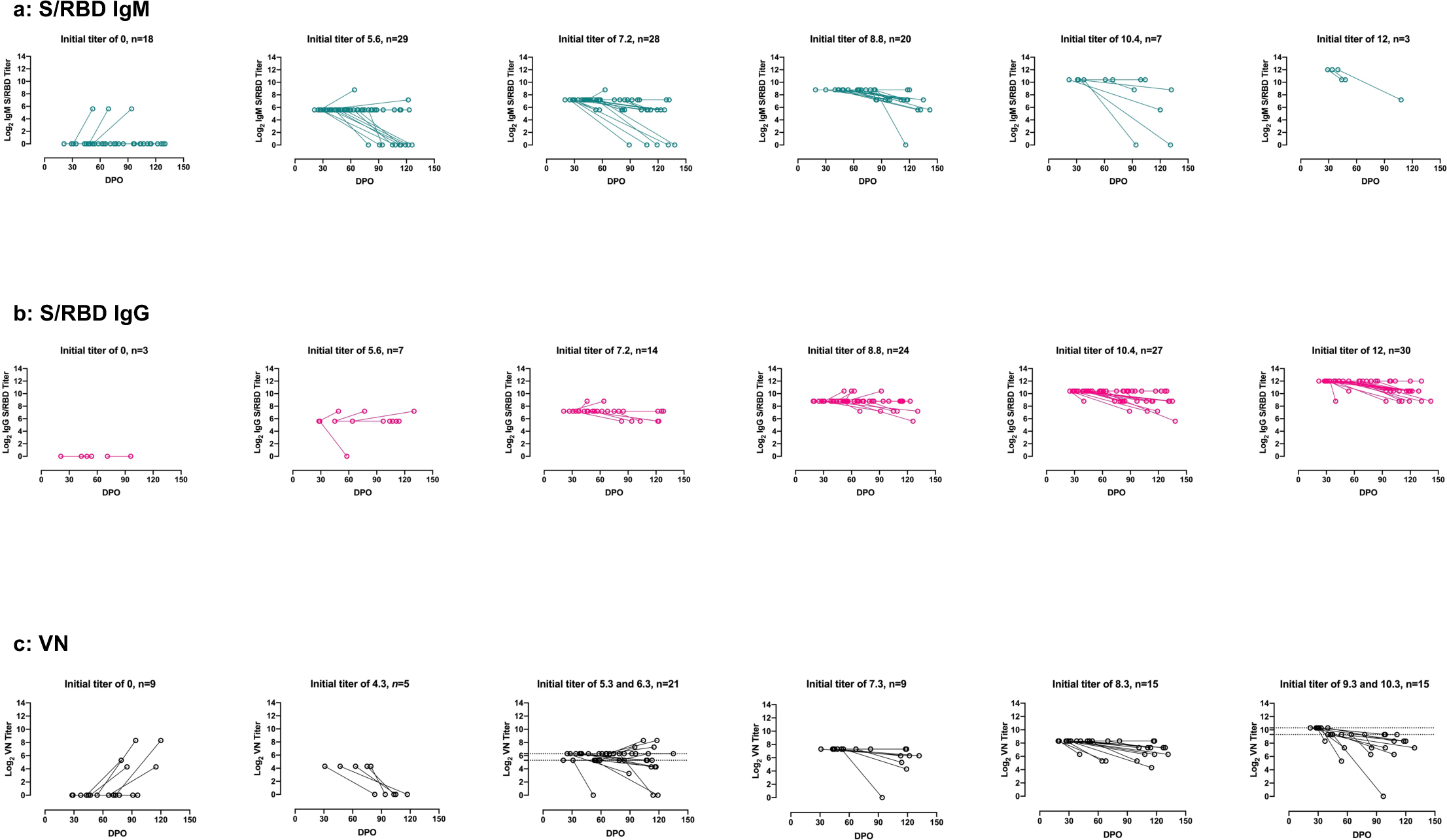
Trajectories (first and last donation only) of (a) SARS-CoV-2 spike-receptor binding domain (S/RBD) IgM, (b) S/RBD IgG, and (c) virus neutralizing (VN) antibody titers against SARS-CoV-2 in subjects who donated plasma more than once. Initial (Log_2_) S/RBD IgM and IgG titers ≥5.3 remain stable or vary by one or two dilutions below or above the initial titer. A majority of individuals (33 out of 39) with initial (Log_2_) VN titers ≥7.3 begin to drop beyond ~60 DPO.

**Extended Data Figure 2.**
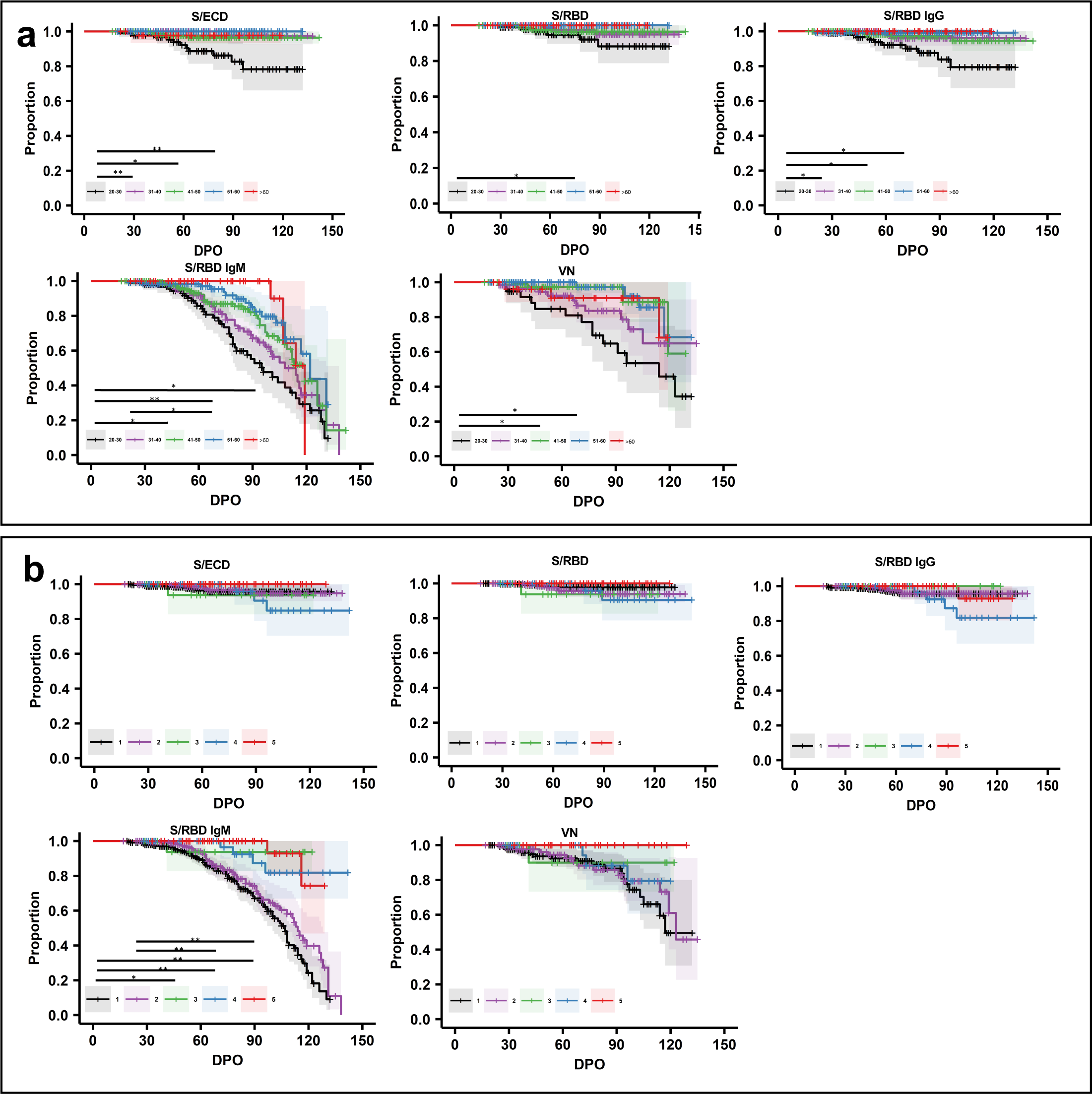
Survival analysis of SARS-CoV-2 spike ectodomain (S/ECD), SARS-CoV-2 spike-receptor binding domain (S/RBD), S/RBD IgM, S/RBD IgG, and neutralizing (VN) antibody titers in 175 convalescent individuals during the first 140 days post onset (DPO) of symptoms stratified by (a) age and (b) severity (Log-rank test, \**P*<0.05, \*\**P*<0.01). Significant differences were observed in the titers of ELISAs between the age groups: 20-30 versus 31-40 (S/ECD \*\**P*<0.01, S/RBD IgG \**P*<0.05); 20-30 versus 41-50 (S/ECD \**P*<0.05, S/RBD IgG \**P*<0.05, S/RBD IgM \**P*<0.05, VN \**P*<0.05); 20-30 versus 51-60 (S/ECD \*\**P*<0.01, S/RBD \**P*<0.05, S/RBD IgG \**P*<0.05, S/RBD IgM \*\**P*<0.01, VN \**P*<0.05); 20-30 versus >60 (S/RBD IgM \**P*<0.05); and 31-40 versus 51-60 (S/RBD IgM \**P*<0.05). Significant differences were observed in the S/RBD IgM titers of the donors with the severity scores 1 versus 3 (\**P*<0.05); 1 versus 4,5 (\*\**P*<0.01); and 2 versus 4,5 (\*\**P*<0.01).

**Extended Data Figure 3.**
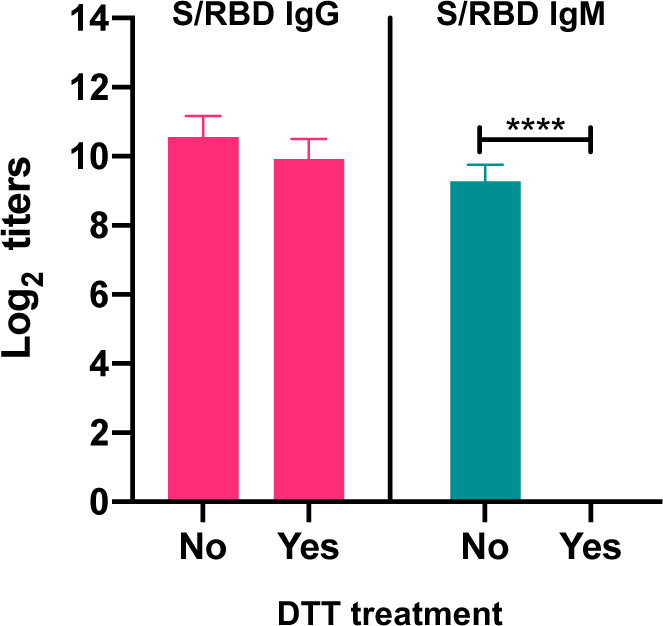
Class specificity test for SARS-CoV-2 spike-receptor binding domain (S/RBD) isotype specific indirect ELISAs. 1,4-Dithiothreitol (DTT) treatment of convalescent plasma abrogates S/RBD IgM antibody titers but not IgG titers (n=10) (paired t test, \*\*\*\**P*<0.0001).

**Extended Data Figure 4.**
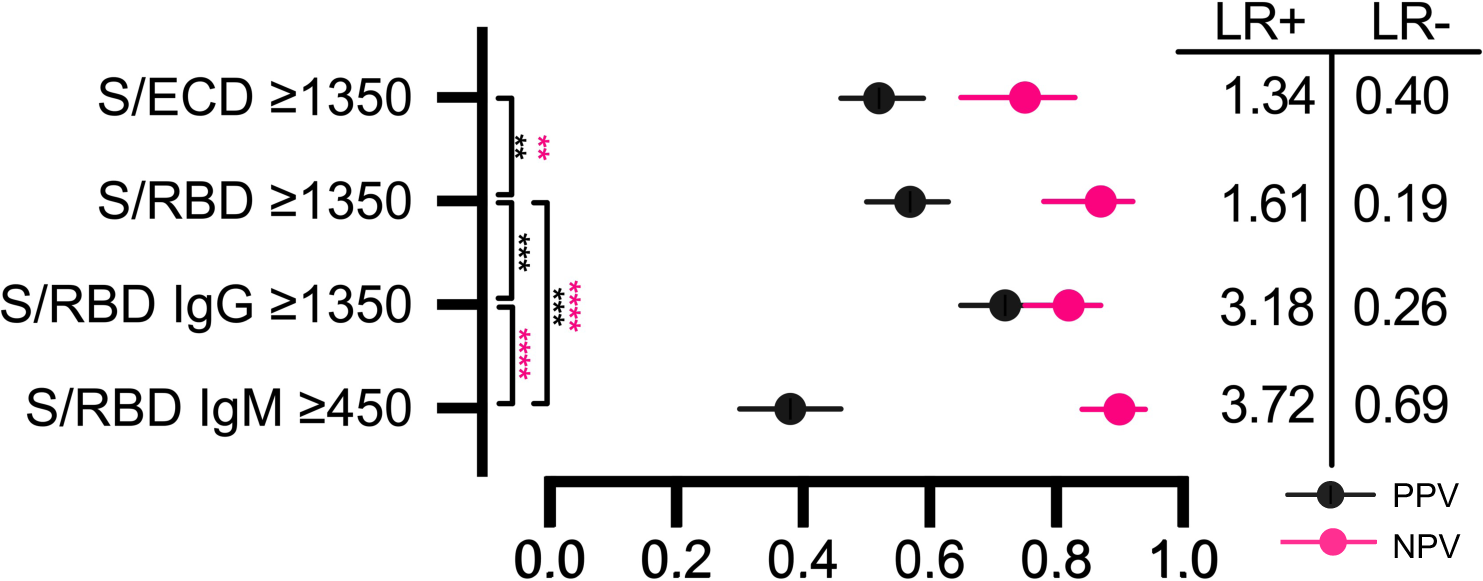
Forest plot depicting the positive and negative predictive values for detection of SARS-CoV-2 spike-receptor binding domain (S/RBD), SARS-CoV-2 spike ectodomain (S/ECD), and S/RBD IgG titers ≥1350 using virus neutralization (VN) titer ≥160 as the standard. Likelihood ratios (LR) for each assay are shown on the right panel. P values were generated using the generalized score statistic for pairwise comparisons. For positive predictive values (PPV) S/ECD ≥1350 versus S/RBD ≥1350 \*\**P*<0.01; S/RBD ≥1350 versus S/RBD IgG ≥1350 \*\*\**P*<0.001; S/RBD IgG ≥1350 versus S/RBD IgM ≥450 P>0.05; S/RBD IgM ≥450 versus S/RBD ≥1350 \*\*\**P*<0.001. For negative predictive values (NPV) S/ECD ≥1350 versus S/RBD ≥1350 \*\**P*<0.01; S/RBD ≥1350 versus S/RBD IgG ≥1350 P>0.05; S/RBD IgG ≥1350 versus S/RBD IgM ≥450 \*\*\*\**P*<0.0001; S/RBD IgM ≥450 versus S/RBD 1350 \*\*\*\**P*<0.0001.

**Supplementary Table 1:** Convalescent plasma donor demographics and sample characteristics.

| Subject | Age | Sex | Hospitalization | Severity | Dyspnea | DPO | VN<br>titer | S/ECD<br>titer | S/RBD<br>titer | S/RBD<br>IgG<br>titer | S/RBD<br>IgM<br>titer |
| --- | --- | --- | --- | --- | --- | --- | --- | --- | --- | --- | --- |
| 0001 | 44 | M | NO | 1 | NO | 20 | 320 | 150 | 1350 | 450 | 150 |
| 0001 |  |  |  |  |  | 24 | 640 | - | - | 450 | 150 |
| 0001 |  |  |  |  |  | 27 | 320 | 150 | 450 | 1350 | 150 |
| 0001 |  |  |  |  |  | 31 | 320 | 150 | 3200 | 1350 | 450 |
| 0001 |  |  |  |  |  | 34 | 320 | 150 | 3200 | 450 | 150 |
| 0001 |  |  |  |  |  | 38 | 160 | 450 | 450 | 450 | 150 |
| 0001 |  |  |  |  |  | 41 | 80 | 450 | 450 | 450 | 150 |
| 0002 | 54 | M | NO | 1 | NO | 28 | 40 | 50 | 150 | 150 | 50 |
| 0003 | 36 | M | NO | 1 | NO | 25 | 80 | 450 | 1350 | 1350 | 150 |
| 0003 |  |  |  |  |  | 28 | 80 | 150 | 450 | 450 | 0 |
| 0003 |  |  |  |  |  | 33 | 80 | 450 | 3200 | 1350 | 50 |
| 0003 |  |  |  |  |  | 35 | 0 | 450 | 450 | 1350 | 50 |
| 0003 |  |  |  |  |  | 42 | 20 | 150 | 450 | 1350 | 50 |
| 0003 |  |  |  |  |  | 47 | 10 | 1350 | 1350 | 450 | 50 |
| 0003 |  |  |  |  |  | 61 | 20 | 1350 | 1350 | 450 | 50 |
| 0003 |  |  |  |  |  | 68 | 0 | 450 | 1350 | 150 | 0 |
| 0003 |  |  |  |  |  | 75 | 40 | 1350 | 450 | 450 | 0 |
| 0003 |  |  |  |  |  | 82 | 20 | 1350 | 450 | 150 | 0 |
| 0003 |  |  |  |  |  | 89 | 10 | 1350 | 1350 | 150 | 0 |
| 0004 | 54 | F | NO | 2 | YES | 32 | 320 | 1350 | 1350 | 1350 | 1350 |
| 0004 |  |  |  |  |  | 36 | 640 | 1350 | 1350 | 4050 | 1350 |
| 0004 |  |  |  |  |  | 68 | 0 | 1350 | 1350 | 1350 | 150 |
| 0004 |  |  |  |  |  | 75 | 80 | 1350 | 1350 | 1350 | 450 |
| 0004 |  |  |  |  |  | 103 | 160 | 1350 | 1350 | 1350 | 150 |
| 0004 |  |  |  |  |  | 118 | 80 | 1350 | 1350 | 1350 | 150 |
| 0004 |  |  |  |  |  | 131 | - | 1350 | 1350 | 450 | 0 |
| 0005 | 58 | M | NO | 2 | YES | 91 | - | 150 | 150 | 150 | 0 |
| 0007 | 36 | M | NO | 1 | NO | 88 | - | 1350 | 1350 | 1350 | 50 |
| 0007 |  |  |  |  |  | 123 | - | 1350 | 1350 | 1350 | 50 |
| 0009 | 38 | F | NO | 2 | YES | 30 | 80 | 450 | 450 | 450 | 50 |
| 0011 | 67 | F | NO | 1 | NO | 28 | 0 | 0 | 50 | 50 | 50 |
| 0011 |  |  |  |  |  | 49 | - | 450 | 150 | 150 | 50 |
| 0012 | 46 | F | NO | 1 | NO | 30 | 320 | 150 | 450 | 150 | 150 |
| 0013 | 43 | F | NO | 1 | NO | 28 | 320 | 1350 | 3200 | 4050 | 450 |
| 0016 | 47 | F | NO | 1 | NO | 32 | 640 | 4050 | 4050 | 4050 | 1350 |
| 0020 | 41 | F | NO | 2 | YES | 17 | 20 | 50 | 200 | 50 | 50 |
| 0022 | 22 | M | NO | 1 | NO | 47 | - | 450 | 150 | 150 | 50 |
| 0022 |  |  |  |  |  | 57 | - | 450 | 1350 | 150 | 50 |
| 0028 | 23 | M | NO | 1 | NO | 31 | 20 | 150 | 150 | 150 | 50 |
| 0028 |  |  |  |  |  | 46 | - | 450 | 150 | 150 | 50 |
| 0028 |  |  |  |  |  | 63 | - | 0 | 450 | 150 | 150 |
| 0028 |  |  |  |  |  | 77 | - | 150 | 150 | 150 | 50 |
| 0028 |  |  |  |  |  | 83 | 0 | 450 | 450 | 50 | 50 |
| 0029 | 66 | F | NO | 1 | NO | 22 | 80 | 150 | 450 | 4050 | 150 |
| 0032 | 65 | M | NO | 2 | YES | 25 | 320 | 450 | 4050 | 4050 | 1350 |
| 0035 | 50 | M | NO | 2 | YES | 28 | 320 | 1350 | 3200 | 4050 | 150 |
| 0035 |  |  |  |  |  | 38 | 640 | 1350 | 1350 | 4050 | 150 |
| 0035 |  |  |  |  |  | 52 | - | 1350 | 1350 | 4050 | 150 |
| 0035 |  |  |  |  |  | 59 | 160 | 1350 | 1350 | 1350 | 50 |
| 0035 |  |  |  |  |  | 82 | 640 | 1350 | 1350 | 1350 | 50 |
| 0035 |  |  |  |  |  | 108 | 80 | 1350 | 1350 | 450 | 0 |
| 0040 | 52 | M | NO | 2 | YES | 29 | 1280 | - | - | 4050 | 4050 |
| 0040 |  |  |  |  |  | 35 | 320 | 4050 | 4050 | 4050 | 4050 |
| 0040 |  |  |  |  |  | 37 | 320 | 1350 | 4050 | 4050 | 4050 |
| 0040 |  |  |  |  |  | 44 | - | 4050 | 4050 | 4050 | 1350 |
| 0045 | 23 | F | NO | 1 | NO | 33 | 1280 | 4050 | 1350 | 4050 | 50 |
| 0045 |  |  |  |  |  | 47 | - | 1350 | 1350 | 4050 | 150 |
| 0045 |  |  |  |  |  | 54 | 40 | 1350 | 1350 | 1350 | 50 |
| 0049 | 57 | F | NO | 1 | NO | 27 | 320 | 150 | 450 | 150 | 50 |
| 0049 |  |  |  |  |  | 64 | 40 | 1350 | 1350 | 450 | 450 |
| 0050 | 41 | M | NO | 2 | YES | 30 | 320 | 1350 | 1350 | 450 | 150 |
| 0050 |  |  |  |  |  | 33 | 0 | 150 | 150 | 1350 | 50 |
| 0050 |  |  |  |  |  | 44 | 20 | 1350 | 450 | 450 | 150 |
| 0050 |  |  |  |  |  | 58 | 20 | 1350 | 1350 | 150 | 50 |
| 0050 |  |  |  |  |  | 66 | 40 | 1350 | 450 | 150 | 50 |
| 0050 |  |  |  |  |  | 72 | 20 | 1350 | 1350 | 150 | 50 |
| 0050 |  |  |  |  |  | 94 | 20 | 1350 | 1350 | 150 | 50 |
| 0050 |  |  |  |  |  | 102 | 20 | 1350 | 1350 | 150 | 50 |
| 0050 |  |  |  |  |  | 108 | - | 1350 | 450 | 450 | 50 |
| 0050 |  |  |  |  |  | 115 | 20 | 1350 | 450 | 150 | 50 |
| 0050 |  |  |  |  |  | 131 | - | 1350 | 1350 | 150 | 0 |
| 0051 | 50 | F | NO | 1 | NO | 30 | 160 | 150 | 450 | 150 | 150 |
| 0052 | 27 | F | YES | 3 | YES | 31 | 160 | 1350 | 1350 | 4050 | 50 |
| 0052 |  |  |  |  |  | 59 | - | 1350 | 1350 | 1350 | 150 |
| 0052 |  |  |  |  |  | 87 | 160 | 1350 | 1350 | 1350 | 150 |
| 0052 |  |  |  |  |  | 122 | 80 | 1350 | 1350 | 1350 | 150 |
| 0053 | 29 | M | NO | 2 | YES | 28 | 0 | 450 | 450 | 450 | 50 |
| 0053 | 30 | M | NO | 2 | YES | 56 | - | 450 | 450 | 450 | 0 |
| 0053 |  |  |  |  |  | 63 | - | 450 | 450 | 150 | 0 |
| 0053 |  |  |  |  |  | 77 | 0 | 1350 | 1350 | 150 | 0 |
| 0053 |  |  |  |  |  | 91 | 0 | 150 | 50 | 150 | 0 |
| 0055 | 61 | M | YES | 3 | YES | 33 | 320 | 1350 | 3200 | 1350 | 150 |
| 0057 | 44 | F | NO | 2 | YES | 34 | 160 | 450 | 450 | 450 | 150 |
| 0058 | 36 | M | NO | 2 | YES | 92 | - | 1350 | 1350 | 1350 | 50 |
| 0062 | 24 | F | NO | 1 | NO | 32 | 320 | 1350 | 1350 | 1350 | 1350 |
| 0062 |  |  |  |  |  | 35 | 1280 | 1350 | 1350 | 1350 | 1350 |
| 0062 |  |  |  |  |  | 46 | - | 1350 | 1350 | 1350 | 1350 |
| 0062 |  |  |  |  |  | 53 | - | 1350 | 1350 | 1350 | 1350 |
| 0062 |  |  |  |  |  | 62 | - | 1350 | 1350 | 1350 | 450 |
| 0062 |  |  |  |  |  | 69 | 320 | 1350 | 1350 | 1350 | 450 |
| 0062 |  |  |  |  |  | 101 | 80 | 1350 | 1350 | 450 | 450 |
| 0062 |  |  |  |  |  | 118 | 160 | 1350 | 1350 | 450 | 450 |
| 0062 |  |  |  |  |  | 132 | 80 | 1350 | 1350 | 450 | 450 |
| 0065 | 50 | M | NO | 1 | NO | 34 | - | 1350 | 1350 | 150 | 0 |
| 0065 |  |  |  |  |  | 54 | - | 1350 | 1350 | 450 | 150 |
| 0065 |  |  |  |  |  | 126 | - | 450 | 450 | 150 | 0 |
| 0069 | 49 | F | NO | 1 | NO | 28 | 80 | 450 | 450 | 1350 | 50 |
| 0069 |  |  |  |  |  | 32 | 80 | 450 | 450 | 1350 | 50 |
| 0069 |  |  |  |  |  | 63 | - | 50 | 450 | 450 | 0 |
| 0069 |  |  |  |  |  | 108 | 40 | 1350 | 450 | 150 | 0 |
| 0070 | 37 | F | NO | 2 | YES | 38 | 160 | 450 | 1350 | 4050 | 50 |
| 0072 | 23 | F | NO | 1 | NO | 29 | 0 | 0 | 0 | 50 | 0 |
| 0072 |  |  |  |  |  | 37 | 0 | 50 | 50 | 0 | 0 |
| 0072 |  |  |  |  |  | 44 | - | 0 | 0 | 0 | 0 |
| 0072 |  |  |  |  |  | 51 | - | 0 | 0 | 0 | 0 |
| 0072 |  |  |  |  |  | 58 | - | 0 | 0 | 0 | 0 |
| 0073 | 39 | F | NO | 2 | YES | 47 | 0 | 150 | 150 | 150 | 0 |
| 0073 |  |  |  |  |  | 55 | - | 450 | 450 | 150 | 0 |
| 0073 |  |  |  |  |  | 85 | 20 | 1350 | 450 | 150 | 0 |
| 0077 | 59 | F | NO | 1 | NO | 71 | - | 1350 | 1350 | 1350 | 150 |
| 0081 | 64 | F | NO | 2 | YES | 52 | - | 450 | 1350 | 1350 | 150 |
| 0088 | 29 | F | NO | 1 | NO | 21 | 20 | 0 | 50 | 0 | 0 |
| 0088 |  |  |  |  |  | 54 | - | 0 | 50 | 0 | 0 |
| 0089 | 42 | F | NO | 2 | YES | 38 | 160 | 450 | 450 | 4050 | 150 |
| 0090 | 33 | M | NO | 1 | NO | 37 | 1280 | 150 | 150 | 150 | 50 |
| 0090 |  |  |  |  |  | 46 | - | 1350 | 1350 | 450 | 50 |
| 0095 | 61 | F | NO | 1 | NO | 54 | 160 | 1350 | 1350 | 4050 | 150 |
| 0095 |  |  |  |  |  | 76 | - | 1350 | 1350 | 1350 | 50 |
| 0095 |  |  |  |  |  | 83 | - | 1350 | 1350 | 1350 | 50 |
| 0095 |  |  |  |  |  | 119 | 20 | 1350 | 1350 | 450 | 0 |
| 0096 | 44 | F | NO | 1 | NO | 59 | - | 1350 | 1350 | 1350 | 50 |
| 0099 | 54 | F | NO | 1 | NO | 20 | 20 | 50 | 50 | 0 | 0 |
| 0109 | 33 | F | NO | 1 | NO | 67 | 160 | 1350 | 1350 | 450 | 50 |
| 0109 |  |  |  |  |  | 73 |  | 1350 | 1350 | 450 | 50 |
| 0109 |  |  |  |  |  | 80 | 160 | 1350 | 1350 | 450 | 0 |
| 0109 |  |  |  |  |  | 93 | 0 | 450 | 150 | 450 | 0 |
| 0109 |  |  |  |  |  | 100 | - | 450 | 450 | 450 | 0 |
| 0109 |  |  |  |  |  | 108 | - | 1350 | 450 | 450 | 0 |
| 0109 |  |  |  |  |  | 114 | 40 | 1350 | 450 | 450 | 0 |
| 0112 | 47 | F | NO | 1 | NO | 32 | 40 | 450 | 450 | 450 | 150 |
| 0113 | 52 | F | NO | 1 | NO | 29 | 40 | 1350 | 150 | 150 | 0 |
| 0115 | 70 | M | YES | 5 | NO | 54 | 320 | 1350 | 1350 | 4050 | 450 |
| 0115 |  |  |  |  |  | 62 | - | 1350 | 1350 | 4050 | 150 |
| 0115 |  |  |  |  |  | 102 | 640 | 1350 | 1350 | 1350 | 150 |
| 0115 |  |  |  |  |  | 118 | 320 | 1350 | 1350 | 1350 | 150 |
| 0116 | 27 | M | NO | 1 | NO | 32 | 20 | 450 | 450 | 450 | 50 |
| 0117 | 27 | F | NO | 2 | YES | 30 | 320 | 1350 | 1350 | 1350 | 150 |
| 0117 |  |  |  |  |  | 44 | - | 1350 | 1350 | 1350 | 150 |
| 0117 |  |  |  |  |  | 71 | - | 1350 | 1350 | 1350 | 150 |
| 0117 |  |  |  |  |  | 79 | 160 | 1350 | 1350 | 1350 | 150 |
| 0117 |  |  |  |  |  | 85 | - | 1350 | 1350 | 1350 | 150 |
| 0117 |  |  |  |  |  | 129 | 160 | 1350 | 1350 | 1350 | 150 |
| 0118 | 50 | F | NO | 1 | NO | 34 | 320 | 1350 | 450 | 4050 | 150 |
| 0118 |  |  |  |  |  | 40 | - | 1350 | 1350 | 450 | 150 |
| 0119 | 35 | F | NO | 1 | NO | 25 | 0 | 450 | 450 | 1350 | 50 |
| 0119 |  |  |  |  |  | 34 | - | 1350 | 1350 | 450 | 50 |
| 0119 |  |  |  |  |  | 40 | - | 1350 | 450 | 450 | 50 |
| 0120 | 41 | F | NO | 1 | NO | 19 | 320 | 450 | 3200 | 450 | 450 |
| 0120 |  |  |  |  |  | 68 | 40 | 1350 | 1350 | 450 | 450 |
| 0121 | 51 | F | NO | 1 | NO | 21 | 40 | 150 | 200 | 150 | 50 |
| 0121 |  |  |  |  |  | 54 | 40 | 450 | 450 | 150 | 50 |
| 0132 | 61 | M | YES | 5 | YES | 78 | 640 | 1350 | 1350 | 4050 | 450 |
| 0132 |  |  |  |  |  | 85 | 160 | 1350 | 1350 | 4050 | 150 |
| 0133 | 51 | M | NO | 2 | YES | 25 | - | 150 | 450 | 450 | 150 |
| 0133 |  |  |  |  |  | 53 | - | 1350 | 1350 | 450 | 50 |
| 0135 | 47 | F | NO | 2 | YES | 32 | 320 | 4050 | 4050 | 4050 | 150 |
| 0137 | 53 | F | NO | 1 | NO | 38 | 80 | 450 | 3200 | 450 | 50 |
| 0137 |  |  |  |  |  | 59 | - | 450 | 1350 | 450 | 50 |
| 0137 |  |  |  |  |  | 66 | - | 1350 | 1350 | 450 | 50 |
| 0137 |  |  |  |  |  | 73 | 80 | 1350 | 1350 | 450 | 50 |
| 0140 | 54 | M | NO | 1 | NO | 98 | - | 1350 | 1350 | 450 | 50 |
| 0143 | 49 | F | NO | 2 | YES | 37 | 160 | 1350 | 3200 | 4050 | 150 |
| 0144 | 48 | M | YES | 5 | YES | 40 | 640 | 1350 | 3200 | 4050 | 450 |
| 0144 |  |  |  |  |  | 45 | 640 | 4050 | 1350 | 4050 | 1350 |
| 0144 |  |  |  |  |  | 50 | 320 | 450 | 1350 | 4050 | 450 |
| 0144 |  |  |  |  |  | 53 | 1280 | 4050 | 4050 | 1350 | 150 |
| 0144 |  |  |  |  |  | 64 | 320 | 1350 | 1350 | 1350 | 150 |
| 0144 |  |  |  |  |  | 73 | 80 | 1350 | 1350 | 1350 | 150 |
| 0144 |  |  |  |  |  | 80 | 160 | 1350 | 1350 | 1350 | 50 |
| 0144 |  |  |  |  |  | 94 | 80 | 1350 | 1350 | 1350 | 150 |
| 0144 |  |  |  |  |  | 115 | 160 | 1350 | 1350 | 1350 | 50 |
| 0144 |  |  |  |  |  | 121 | - | 1350 | 450 | 1350 | 50 |
| 0144 |  |  |  |  |  | 129 | 160 | 1350 | 1350 | 1350 | 50 |
| 0156 | 59 | M | NO | 1 | NO | 22 | 1280 | 4050 | 1350 | 4050 | 1350 |
| 0156 |  |  |  |  |  | 29 | 1280 | 1350 | 1350 | 4050 | 450 |
| 0156 |  |  |  |  |  | 43 | 80 | 1350 | 1350 | 4050 | 450 |
| 0156 |  |  |  |  |  | 50 | - | 1350 | 1350 | 4050 | 450 |
| 0156 |  |  |  |  |  | 65 | - | 1350 | 1350 | 4050 | 150 |
| 0156 |  |  |  |  |  | 71 | 160 | 1350 | 1350 | 1350 | 150 |
| 0156 |  |  |  |  |  | 85 | - | 1350 | 1350 | 1350 | 50 |
| 0156 |  |  |  |  |  | 92 | - | 1350 | 1350 | 1350 | 50 |
| 0156 |  |  |  |  |  | 99 | 160 | 1350 | 1350 | 1350 | 50 |
| 0156 |  |  |  |  |  | 106 | - | 1350 | 1350 | 1350 | 50 |
| 0156 |  |  |  |  |  | 113 | - | 1350 | 1350 | 1350 | 50 |
| 0156 |  |  |  |  |  | 120 | 320 | 1350 | 1350 | 1350 | 50 |
| 0158 | 33 | M | NO | 2 | YES | 26 | 40 | 150 | 200 | 450 | 50 |
| 0159 | 23 | F | NO | 2 | YES | 46 | - | 4050 | 4050 | 4050 | 450 |
| 0162 | 51 | F | YES | 3 | YES | 34 | 160 | 150 | 450 | 450 | 50 |
| 0177 | 55 | M | NO | 1 | NO | 44 | 160 | 4050 | 4050 | 4050 | 450 |
| 0177 |  |  |  |  |  | 132 | 80 | 1350 | 1350 | 450 | 50 |
| 0215 | 38 | M | NO | 2 | YES | 53 | - | 1350 | 1350 | 450 | 50 |
| 0215 |  |  |  |  |  | 60 | - | 1350 | 1350 | 1350 | 50 |
| 0229 | 32 | M | NO | 2 | YES | 40 | 80 | 450 | 1350 | 1350 | 450 |
| 0229 |  |  |  |  |  | 61 | 320 | 1350 | 1350 | 1350 | 450 |
| 0229 |  |  |  |  |  | 68 | - | 1350 | 1350 | 450 | 150 |
| 0229 |  |  |  |  |  | 76 | - | 1350 | 1350 | 450 | 150 |
| 0229 |  |  |  |  |  | 100 | 160 | 1350 | 1350 | 450 | 150 |
| 0229 |  |  |  |  |  | 110 | - | 1350 | 1350 | 450 | 150 |
| 0229 |  |  |  |  |  | 117 | 160 | 1350 | 1350 | 450 | 50 |
| 0229 |  |  |  |  |  | 135 | 80 | 1350 | 1350 | 450 | 150 |
| 0234 | 40 | M | NO | 2 | YES | 27 | 40 | 150 | 150 | 150 | 150 |
| 0245 | 51 | M | YES | 5 | YES | 38 | 320 | 1350 | 4050 | 4050 | 150 |
| 0245 |  |  |  |  |  | 52 | - | 1350 | 1350 | 4050 | 450 |
| 0245 |  |  |  |  |  | 59 | - | 1350 | 1350 | 4050 | 150 |
| 0245 |  |  |  |  |  | 81 | 320 | 1350 | 1350 | 1350 | 150 |
| 0245 |  |  |  |  |  | 102 | 160 | 1350 | 1350 | 1350 | 50 |
| 0249 | 56 | M | NO | 1 | NO | 22 | 320 | 4050 | 4050 | 4050 | 150 |
| 0255 | 40 | M | NO | 2 | YES | 31 | 40 | 450 | 450 | 450 | 0 |
| 0255 |  |  |  |  |  | 45 | - | 1350 | 1350 | 1350 | 50 |
| 0255 |  |  |  |  |  | 52 | 0 | 1350 | 1350 | 1350 | 50 |
| 0260 | 44 | M | NO | 2 | YES | 24 | 1280 | 4050 | 1350 | 1350 | 1350 |
| 0262 | 36 | F | YES | 4 | YES | 31 | 1280 | 4050 | 4050 | 4050 | 1350 |
| 0262 |  |  |  |  |  | 49 | - | 1350 | 1350 | 4050 | 1350 |
| 0262 |  |  |  |  |  | 99 | 640 | 1350 | 1350 | 4050 | 1350 |
| 0263 | 20 | M | NO | 1 | NO | 43 | 0 | 1350 | 1350 | 1350 | 50 |
| 0263 |  |  |  |  |  | 52 | - | 150 | 1350 | 450 | 50 |
| 0263 |  |  |  |  |  | 59 | - | 1350 | 450 | 450 | 0 |
| 0263 |  |  |  |  |  | 79 | 40 | 1350 | 1350 | 450 | 0 |
| 0265 | 53 | F | NO | 2 | YES | 31 | 320 | 4050 | 4050 | 4050 | 150 |
| 0280 | 37 | M | YES | 4 | YES | 73 | 0 | 1350 | 1350 | 4050 | 450 |
| 0280 |  |  |  |  |  | 98 | 320 | 1350 | 1350 | 4050 | 450 |
| 0280 |  |  |  |  |  | 120 | 320 | 1350 | 1350 | 4050 | 450 |
| 0284 | 35 | F | NO | 4 | YES | 53 | - | 1350 | 1350 | 1350 | 150 |
| 0285 | 51 | F | NO | 1 | NO | 40 | - | 1350 | 1350 | 450 | 150 |
| 0285 |  |  |  |  |  | 56 | - | 1350 | 1350 | 450 | 150 |
| 0287 | 40 | F | NO | 2 | YES | 56 | - | 50 | 0 | 0 | 0 |
| 0301 | 59 | M | NO | 1 | NO | 83 | - | 1350 | 1350 | 1350 | 450 |
| 0301 |  |  |  |  |  | 90 | - | 1350 | 1350 | 1350 | 450 |
| 0301 |  |  |  |  |  | 97 | - | 1350 | 1350 | 450 | 150 |
| 0302 | 25 | M | NO | 1 | NO | 84 | 80 | 1350 | 1350 | 450 | 150 |
| 0302 |  |  |  |  |  | 96 | 80 | 450 | 150 | 450 | 50 |
| 0302 |  |  |  |  |  | 112 | - | 450 | 1350 | 450 | 50 |
| 0313 | 78 | M | YES | 3 | YES | 36 | 160 | 4050 | 4050 | 4050 | 50 |
| 0339 | 31 | M | YES | 3 | YES | 54 | 640 | 150 | 1350 | 4050 | 450 |
| 0339 |  |  |  |  |  | 62 | - | 1350 | 1350 | 4050 | 450 |
| 0339 |  |  |  |  |  | 68 | 160 | 1350 | 1350 | 4050 | 450 |
| 0339 |  |  |  |  |  | 82 | - | 1350 | 1350 | 1350 | 150 |
| 0339 |  |  |  |  |  | 89 | - | 1350 | 1350 | 1350 | 150 |
| 0339 |  |  |  |  |  | 96 | 160 | 1350 | 1350 | 1350 | 150 |
| 0339 |  |  |  |  |  | 110 | - | 1350 | 1350 | 1350 | 150 |
| 0339 |  |  |  |  |  | 118 | 320 | 1350 | 1350 | 1350 | 150 |
| 0345 | 62 | M | NO | 1 | NO | 54 | 0 | 1350 | 1350 | 4050 | 450 |
| 0345 |  |  |  |  |  | 94 | 320 | 1350 | 1350 | 1350 | 150 |
| 0350 | 53 | M | NO | 1 | NO | 45 | 160 | 1350 | 1350 | 1350 | 150 |
| 0350 |  |  |  |  |  | 59 | - | 1350 | 1350 | 1350 | 150 |
| 0350 |  |  |  |  |  | 82 | 160 | 1350 | 1350 | 450 | 50 |
| 0354 | 59 | M | YES | 4 | YES | 98 | 640 | 1350 | 1350 | 4050 | 150 |
| 0354 |  |  |  |  |  | 104 | - | 1350 | 1350 | 4050 | 150 |
| 0354 |  |  |  |  |  | 111 | 640 | 1350 | 1350 | 4050 | 450 |
| 0354 |  |  |  |  |  | 132 | - | 1350 | 1350 | 4050 | 150 |
| 0363 | 56 | M | NO | 1 | NO | 34 | 80 | 450 | 1350 | 1350 | 50 |
| 0363 |  |  |  |  |  | 44 | - | 1350 | 1350 | 1350 | 50 |
| 0363 |  |  |  |  |  | 59 | - | 1350 | 1350 | 450 | 50 |
| 0363 |  |  |  |  |  | 64 | - | 1350 | 1350 | 450 | 50 |
| 0363 |  |  |  |  |  | 73 | 80 | 1350 | 1350 | 450 | 50 |
| 0363 |  |  |  |  |  | 86 | - | 450 | 1350 | 450 | 0 |
| 0363 |  |  |  |  |  | 100 | 20 | 1350 | 1350 | 450 | 50 |
| 0363 |  |  |  |  |  | 107 | 40 | 1350 | 450 | 450 | 50 |
| 0367 | 58 | F | NO | 2 | YES | 89 | - | 1350 | 1350 | 4050 | 450 |
| 0368 | 37 | F | NO | 2 | YES | 29 | 160 | 450 | 1350 | 4050 | 50 |
| 0369 | 41 | M | NO | 1 | NO | 39 | 80 | 150 | 450 | 1350 | 50 |
| 0369 |  |  |  |  |  | 46 | 40 | 450 | 450 | 450 | 50 |
| 0369 |  |  |  |  |  | 49 | 40 | 1350 | 1350 | 450 | 50 |
| 0369 |  |  |  |  |  | 56 | 80 | 1350 | 1350 | 450 | 50 |
| 0369 |  |  |  |  |  | 63 | 20 | 1350 | 1350 | 450 | 0 |
| 0369 |  |  |  |  |  | 69 | 20 | 1350 | 450 | 450 | 50 |
| 0369 |  |  |  |  |  | 76 | 20 | 1350 | 450 | 150 | 50 |
| 0369 |  |  |  |  |  | 83 | 10 | 1350 | 1350 | 150 | 50 |
| 0369 |  |  |  |  |  | 97 | 20 | 450 | 450 | 450 | 0 |
| 0369 |  |  |  |  |  | 104 | - | 450 | 150 | 150 | 0 |
| 0369 |  |  |  |  |  | 112 | 20 | 1350 | 450 | 150 | 0 |
| 0369 | 41 | M | NO | 1 | NO | 119 | - | 450 | 450 | 150 | 0 |
| 0376 | 52 | M | YES | 4 | YES | 28 | 1280 | 1350 | 4050 | 4050 | 150 |
| 0376 |  |  |  |  |  | 32 | 160 | 4050 | 4050 | 4050 | 150 |
| 0376 |  |  |  |  |  | 60 | - | 1350 | 1350 | 1350 | 150 |
| 0376 |  |  |  |  |  | 67 | 80 | 1350 | 1350 | 1350 | 50 |
| 0376 |  |  |  |  |  | 88 | 160 | 1350 | 1350 | 1350 | 50 |
| 0376 |  |  |  |  |  | 108 | 80 | 1350 | 1350 | 450 | 50 |
| 0377 | 49 | M | NO | 2 | YES | 52 | 160 | 450 | 1350 | 150 | 0 |
| 0377 |  |  |  |  |  | 66 | - | 50 | 450 | 150 | 0 |
| 0377 |  |  |  |  |  | 80 | - | 450 | 450 | 50 | 0 |
| 0377 |  |  |  |  |  | 94 | 0 | 450 | 450 | 50 | 50 |
| 0385 | 45 | M | NO | 2 | YES | 63 | - | 0 | 0 | 0 | 0 |
| 0398 | 55 | F | YES | 4 | YES | 58 | - | 1350 | 1350 | 1350 | 150 |
| 0398 |  |  |  |  |  | 67 | - | 1350 | 1350 | 1350 | 150 |
| 0398 |  |  |  |  |  | 73 | - | 1350 | 1350 | 450 | 150 |
| 0412 | 53 | M | NO | 1 | NO | 75 | 20 | 1350 | 450 | 150 | 0 |
| 0412 |  |  |  |  |  | 89 | - | 1350 | 1350 | 150 | 0 |
| 0412 |  |  |  |  |  | 103 | 0 | 1350 | 450 | 50 | 0 |
| 0419 | 40 | M | NO | 2 | YES | 68 | -- | 1350 | 1350 | 1350 | 150 |
| 0422 | 41 | M | NO | 1 | NO | 64 | 640 | 1350 | 1350 | 1350 | 450 |
| 0422 |  |  |  |  |  | 84 | 80 | 1350 | 450 | 450 | 150 |
| 0423 | 39 | M | NO | 2 | YES | 57 | 40 | 450 | 1350 | 1350 | 150 |
| 0423 |  |  |  |  |  | 65 | 20 | 1350 | 1350 | 450 | 50 |
| 0423 |  |  |  |  |  | 75 | 20 | 50 | 1350 | 450 | 50 |
| 0423 |  |  |  |  |  | 83 | 20 | 1350 | 1350 | 450 | 50 |
| 0423 |  |  |  |  |  | 89 | 20 | 450 | 1350 | 150 | 50 |
| 0423 |  |  |  |  |  | 117 | 20 | 450 | 450 | 150 | 0 |
| 0423 |  |  |  |  |  | 127 | - | 1350 | 450 | 150 | 0 |
| 0423 |  |  |  |  |  | 131 | - | 1350 | 1350 | 150 | 0 |
| 0423 |  |  |  |  |  | 138 | - | 450 | 1350 | 50 | 0 |
| 0430 | 44 | M | YES | 4 | YES | 35 | 1280 | 4050 | 4050 | 4050 | 150 |
| 0436 | 32 | F | NO | 1 | NO | 45 | 640 | 1350 | 1350 | 50 | 0 |
| 0436 |  |  |  |  |  | 55 | - | 1350 | 1350 | 50 | 0 |
| 0436 |  |  |  |  |  | 62 | - | 450 | 1350 | 50 | 0 |
| 0436 |  |  |  |  |  | 69 | - | 1350 | 1350 | 50 | 0 |
| 0436 |  |  |  |  |  | 90 | - | 1350 | 1350 | 50 | 0 |
| 0436 |  |  |  |  |  | 97 | 0 | 1350 | 1350 | 50 | 0 |
| 0437 | 49 | M | NO | 1 | NO | 55 | 40 | 1350 | 1350 | 450 | 50 |
| 0437 |  |  |  |  |  | 64 | - | 1350 | 1350 | 450 | 50 |
| 0437 |  |  |  |  |  | 78 | - | 1350 | 1350 | 450 | 50 |
| 0437 |  |  |  |  |  | 113 | 40 | 1350 | 1350 | 450 | 50 |
| 0448 | 49 | M | NO | 2 | YES | 43 | 160 | 1350 | 1350 | 4050 | 150 |
| 0448 |  |  |  |  |  | 46 | - | 1350 | 1350 | 4050 | 150 |
| 0448 |  |  |  |  |  | 91 | 160 | 1350 | 1350 | 1350 | 150 |
| 0448 |  |  |  |  |  | 105 | - | 1350 | 1350 | 4050 | 150 |
| 0448 |  |  |  |  |  | 112 | - | 1350 | 1350 | 1350 | 50 |
| 0448 |  |  |  |  |  | 119 | 160 | 1350 | 1350 | 1350 | 50 |
| 0462 | 47 | F | NO | 2 | YES | 61 | - | 450 | 1350 | 450 | 0 |
| 0464 | 31 | F | NO | 2 | YES | 48 | 160 | 1350 | 1350 | 4050 | 450 |
| 0464 |  |  |  |  |  | 55 | 160 | 1350 | 1350 | 4050 | 450 |
| 0464 |  |  |  |  |  | 62 | 80 | 1350 | 1350 | 450 | 450 |
| 0464 |  |  |  |  |  | 69 | 80 | 1350 | 1350 | 1350 | 450 |
| 0464 |  |  |  |  |  | 83 | 80 | 1350 | 1350 | 1350 | 150 |
| 0464 |  |  |  |  |  | 90 | - | 1350 | 1350 | 1350 | 450 |
| 0464 |  |  |  |  |  | 118 | 160 | 1350 | 1350 | 1350 | 450 |
| 0479 | 56 | F | NO | 1 | NO | 79 | - | 1350 | 1350 | 4050 | 450 |
| 0488 | 37 | F | NO | 1 | NO | 62 | - | 150 | 150 | 0 | 0 |
| 0515 | 58 | M | YES | 4 | YES | 69 | 80 | 1350 | 1350 | 4050 | 1350 |
| 0515 |  |  |  |  |  | 83 | - | 1350 | 1350 | 4050 | 1350 |
| 0515 |  |  |  |  |  | 104 | 320 | 1350 | 1350 | 4050 | 1350 |
| 0524 | 35 | F | YES | 3 | YES | 44 | - | 1350 | 1350 | 4050 | 450 |
| 0525 | 33 | F | NO | 2 | YES | 43 | - | 450 | 50 | 150 | 50 |
| 0525 |  |  |  |  |  | 68 | - | 50 | 450 | 150 | 50 |
| 0526 | 74 | M | NO | 2 | YES | 45 | - | 1350 | 1350 | 1350 | 150 |
| 0530 | 39 | F | NO | 1 | NO | 71 | - | 1350 | 1350 | 450 | 50 |
| 0533 | 32 | F | NO | 1 | NO | 47 | - | 450 | 450 | 150 | 0 |
| 0548 | 40 | M | NO | 4 | NO | 52 | - | 1350 | 1350 | 4050 | 150 |
| 0554 | 68 | F | NO | 1 | NO | 57 | - | 1350 | 1350 | 4050 | 1350 |
| 0576 | 50 | F | YES | 3 | YES | 41 | 0 | 0 | 0 | 50 | 0 |
| 0579 | 70 | M | NO | 2 | YES | 43 | 640 | 1350 | 1350 | 4050 | 450 |
| 0579 |  |  |  |  |  | 50 | - | 1350 | 1350 | 1350 | 150 |
| 0579 |  |  |  |  |  | 57 | - | 1350 | 1350 | 4050 | 450 |
| 0579 |  |  |  |  |  | 64 | 80 | 1350 | 1350 | 1350 | 150 |
| 0579 |  |  |  |  |  | 99 | 160 | 1350 | 1350 | 1350 | 150 |
| 0580 | 43 | F | YES | 3 | YES | 29 | 1280 | 4050 | 4050 | 1350 | 150 |
| 0580 |  |  |  |  |  | 35 | 1280 | 4050 | 4050 | 4050 | 150 |
| 0580 |  |  |  |  |  | 57 | 160 | 1350 | 1350 | 1350 | 50 |
| 0581 | 50 | M | YES | 4 | YES | 108 | - | 1350 | 1350 | 150 | 50 |
| 0591 | 51 | F | NO | 1 | NO | 44 | - | 450 | 1350 | 450 | 150 |
| 0595 | 26 | F | NO | 2 | YES | 104 | - | 150 | 150 | 50 | 0 |
| 0595 |  |  |  |  |  | 111 | - | 150 | 150 | 50 | 0 |
| 0598 | 46 | M | YES | 4 | YES | 30 | 1280 | 4050 | 4050 | 4050 | 450 |
| 0598 |  |  |  |  |  | 99 | 640 | 1350 | 1350 | 4050 | 150 |
| 0599 | 42 | M | YES | 3 | YES | 82 | 320 | 1350 | 1350 | 1350 | 450 |
| 0599 |  |  |  |  |  | 117 | 320 | 1350 | 1350 | 1350 | 150 |
| 0605 | 54 | F | NO | 1 | NO | 58 | - | 1350 | 1350 | 1350 | 150 |
| 0610 | 48 | F | NO | 2 | YES | 64 | - | 1350 | 1350 | 150 | 0 |
| 0612 | 36 | F | NO | 1 | NO | 69 | - | 1350 | 1350 | 50 | 0 |
| 0618 | 58 | F | YES | 5 | YES | 65 | 80 | 1350 | 1350 | 4050 | 450 |
| 0618 |  |  |  |  |  | 118 | 320 | 1350 | 1350 | 1350 | 150 |
| 0620 | 59 | M | YES | 5 | YES | 40 | 1280 | 4050 | 4050 | 4050 | 4050 |
| 0620 |  |  |  |  |  | 52 | 320 | 1350 | 1350 | 4050 | 1350 |
| 0620 |  |  |  |  |  | 59 | 320 | 1350 | 1350 | 4050 | 1350 |
| 0620 |  |  |  |  |  | 66 | 160 | 1350 | 1350 | 4050 | 1350 |
| 0620 |  |  |  |  |  | 73 | 160 | 1350 | 1350 | 4050 | 450 |
| 0620 |  |  |  |  |  | 94 | 320 | 1350 | 1350 | 450 | 450 |
| 0620 |  |  |  |  |  | 101 | -- | 1350 | 1350 | 4050 | 150 |
| 0620 |  |  |  |  |  | 108 | 320 | 1350 | 1350 | 1350 | 150 |
| 0622 | 20 | F | NO | 1 | NO | 45 | 0 | 450 | 150 | 50 | 0 |
| 0622 |  |  |  |  |  | 52 | - | 450 | 150 | 50 | 0 |
| 0622 |  |  |  |  |  | 77 | 0 | 1350 | 450 | 150 | 0 |
| 0631 | 58 | M | NO | 1 | NO | 47 | 20 | 150 | 450 | 150 | 50 |
| 0631 |  |  |  |  |  | 53 | 20 | 150 | 1350 | 50 | 50 |
| 0631 |  |  |  |  |  | 61 | 20 | 1350 | 1350 | 50 | 0 |
| 0631 |  |  |  |  |  | 67 | 20 | 1350 | 150 | 50 | 0 |
| 0631 |  |  |  |  |  | 74 | 10 | 450 | 1350 | 50 | 0 |
| 0631 |  |  |  |  |  | 81 | 20 | 450 | 450 | 50 | 0 |
| 0631 |  |  |  |  |  | 95 | 0 | 450 | 450 | 50 | 0 |
| 0631 |  |  |  |  |  | 108 | - | 150 | 50 | 50 | 0 |
| 0631 |  |  |  |  |  | 117 | 0 | 450 | 450 | 50 | 0 |
| 0631 |  |  |  |  |  | 122 | - | 450 | 150 | 50 | 0 |
| 0633 | 63 | M | YES | 4 | YES | 79 | - | 1350 | 1350 | 1350 | 150 |
| 0634 | 53 | M | YES | 5 | YES | 33 | 1280 | 4050 | 4050 | 4050 | 450 |
| 0636 | 30 | M | NO | 2 | YES | 64 | - | 1350 | 1350 | 1350 | 450 |
| 0664 | 46 | M | NO | 1 | NO | 75 | - | 1350 | 1350 | 1350 | 50 |
| 0664 |  |  |  |  |  | 86 | - | 1350 | 1350 | 1350 | 50 |
| 0694 | 30 | M | NO | 2 | YES | 42 | 320 | 1350 | 1350 | 1350 | 150 |
| 0694 |  |  |  |  |  | 127 | 160 | 1350 | 1350 | 1350 | 50 |
| 0695 | 22 | F | NO | 2 | YES | 62 | 0 | 0 | 1350 | 150 | 0 |
| 0695 |  |  |  |  |  | 72 | - | 450 | 1350 | 150 | 0 |
| 0695 |  |  |  |  |  | 79 | - | 450 | 450 | 150 | 0 |
| 0695 |  |  |  |  |  | 114 | 0 | 450 | 150 | 150 | 0 |
| 0695 |  |  |  |  |  | 128 | - | 450 | 450 | 150 | 0 |
| 0698 | 32 | M | NO | 2 | YES | 51 | 320 | 1350 | 1350 | 450 | 150 |
| 0698 |  |  |  |  |  | 59 | - | 1350 | 1350 | 450 | 150 |
| 0698 |  |  |  |  |  | 72 | - | 1350 | 450 | 450 | 50 |
| 0698 |  |  |  |  |  | 100 | 40 | 1350 | 150 | 450 | 150 |
| 0699 | 61 | M | YES | 3 | NO | 35 | 640 | 4050 | 450 | 450 | 450 |
| 0701 | 41 | M | NO | 1 | NO | 47 | 80 | 1350 | 1350 | 450 | 150 |
| 0701 |  |  |  |  |  | 53 | 40 | 1350 | 1350 | 1350 | 150 |
| 0701 |  |  |  |  |  | 63 | 40 | 1350 | 1350 | 450 | 150 |
| 0701 |  |  |  |  |  | 70 | 40 | 1350 | 1350 | 450 | 150 |
| 0701 |  |  |  |  |  | 74 | 80 | 1350 | 1350 | 150 | 150 |
| 0701 |  |  |  |  |  | 105 | 20 | 1350 | 450 | 450 | 50 |
| 0701 |  |  |  |  |  | 109 | - | 1350 | 1350 | 50 | 0 |
| 0701 |  |  |  |  |  | 116 | 20 | 1350 | 450 | 450 | 50 |
| 0701 |  |  |  |  |  | 123 | - | 1350 | 1350 | 450 | 50 |
| 0719 | 63 | M | NO | 1 | NO | 107 | - | 1350 | 1350 | 50 | 0 |
| 0719 |  |  |  |  |  | 114 | - | 1350 | 1350 | 50 | 0 |
| 0720 | 39 | M | NO | 2 | YES | 63 | - | 50 | 0 | 0 | 0 |
| 0731 | 50 | M | YES | 4 | YES | 66 | 80 | 1350 | 1350 | 4050 | 450 |
| 0731 |  |  |  |  |  | 73 | 40 | 1350 | 1350 | 4050 | 150 |
| 0731 |  |  |  |  |  | 80 | 40 | 1350 | 1350 | 1350 | 50 |
| 0731 |  |  |  |  |  | 87 | 80 | 1350 | 1350 | 450 | 150 |
| 0731 |  |  |  |  |  | 94 | 40 | 1350 | 1350 | 1350 | 150 |
| 0731 |  |  |  |  |  | 101 | - | 1350 | 1350 | 4050 | 150 |
| 0731 |  |  |  |  |  | 115 | 160 | 1350 | 1350 | 1350 | 150 |
| 0731 |  |  |  |  |  | 128 | - | 1350 | 1350 | 1350 | 150 |
| 0731 |  |  |  |  |  | 142 | - | 1350 | 1350 | 450 | 50 |
| 0749 | 45 | M | NO | 1 | NO | 48 | - | 450 | 1350 | 450 | 0 |
| 0749 |  |  |  |  |  | 55 | - | 450 | 1350 | 450 | 0 |
| 0749 |  |  |  |  |  | 69 | - | 450 | 450 | 150 | 50 |
| 0750 | 29 | M | NO | 2 | YES | 46 | - | 450 | 150 | 150 | 150 |
| 0750 |  |  |  |  |  | 50 | - | 450 | 150 | 450 | 50 |
| 0750 |  |  |  |  |  | 53 | - | 150 | 1350 | 150 | 50 |
| 0750 |  |  |  |  |  | 123 | 0 | 450 | 450 | 50 | 50 |
| 0759 | 48 | F | NO | 2 | YES | 45 | - | 1350 | 1350 | 450 | 150 |
| 0762 | 47 | M | NO | 2 | YES | 44 | 160 | 450 | 1350 | 1350 | 50 |
| 0762 |  |  |  |  |  | 51 | 40 | 1350 | 1350 | 1350 | 150 |
| 0762 |  |  |  |  |  | 58 | 80 | 1350 | 1350 | 450 | 150 |
| 0762 |  |  |  |  |  | 64 | 160 | 1350 | 1350 | 450 | 150 |
| 0762 |  |  |  |  |  | 78 | 160 | 1350 | 1350 | 1350 | 50 |
| 0762 |  |  |  |  |  | 85 | - | 1350 | 1350 | 150 | 0 |
| 0762 |  |  |  |  |  | 92 | 40 | 1350 | 1350 | 450 | 0 |
| 0762 |  |  |  |  |  | 113 | 80 | 1350 | 1350 | 450 | 0 |
| 0786 | 46 | F | NO | 2 | YES | 48 | - | 450 | 1350 | 1350 | 150 |
| 0789 | 55 | F | NO | 2 | YES | 62 | - | 1350 | 1350 | 1350 | 150 |
| 0796 | 45 | M | YES | 5 | YES | 52 | - | 1350 | 1350 | 1350 | 450 |
| 0820 | 56 | F | NO | 1 | NO | 82 | - | 1350 | 1350 | 450 | 50 |
| 0820 |  |  |  |  |  | 113 | - | 1350 | 1350 | 450 | 50 |
| 0834 | 48 | M | NO | 2 | YES | 60 | - | 1350 | 1350 | 4050 | 450 |
| 0835 | 52 | F | YES | 5 | YES | 61 | - | 1350 | 1350 | 4050 | 4050 |
| 0838 | 74 | F | NO | 1 | NO | 49 | 320 | 1350 | 1350 | 1350 | 450 |
| 0838 |  |  |  |  |  |  | 160 | 1350 | 1350 | 1350 | 450 |
| 0838 |  |  |  |  |  |  | 160 | 1350 | 1350 | 450 | 450 |
| 0838 |  |  |  |  |  |  | 80 | 1350 | 1350 | 1350 | 450 |
| 0838 |  |  |  |  |  |  | 320 | 1350 | 1350 | 1350 | 150 |
| 0838 |  |  |  |  |  |  | - | 1350 | 1350 | 1350 | 150 |
| 0838 |  |  |  |  |  |  | 160 | 1350 | 1350 | 450 | 150 |
| 0850 | 34 | F | NO | 2 | YES | 67 | - | 1350 | 1350 | 1350 | 150 |
| 0879 | 50 | M | YES | 4 | YES | 34 | 160 | 4050 | 4050 | 4050 | 4050 |
| 0879 |  |  |  |  |  |  | - | 1350 | 1350 | 4050 | 1350 |
| 0905 | 41 | F | NO | 1 | NO | 57 | - | 1350 | 1350 | 150 | 0 |
| 0913 | 34 | F | YES | 5 | YES | 83 | - | 1350 | 1350 | 4050 | 450 |
| 0913 |  |  |  |  |  | 90 | - | 1350 | 1350 | 4050 | 150 |
| 0913 |  |  |  |  |  | 101 | - | 1350 | 1350 | 4050 | 150 |
| 0913 |  |  |  |  |  | 116 | - | 1350 | 1350 | 1350 | 0 |
| 0933 | 67 | M | NO | 1 | NO | 65 | - | 1350 | 1350 | 4050 | 450 |
| 0970 | 29 | F | NO | 2 | YES | 56 | - | 1350 | 1350 | 1350 | 50 |
| 0992 | 45 | F | NO | 2 | YES | 43 | - | 0 | 0 | 0 | 0 |
| 0992 |  |  |  |  |  | 49 | - | 0 | 0 | 0 | 0 |
| 1033 | 33 | F | NO | 1 | NO | 61 | 80 | 1350 | 1350 | 1350 | 1350 |
| 1033 |  |  |  |  |  | 75 | - | 1350 | 450 | 150 | 0 |
| 1033 |  |  |  |  |  | 92 | 80 | 1350 | 1350 | 1350 | 450 |
| 1052 | 28 | F | NO | 2 | YES | 59 | 40 | 1350 | 1350 | 1350 | 50 |
| 1052 |  |  |  |  |  | 73 | - | 150 | 1350 | 450 | 50 |
| 1052 |  |  |  |  |  | 95 | 160 | 1350 | 1350 | 1350 | 50 |
| 1062 | 44 | F | NO | 2 | YES | 53 | 40 | 1350 | 1350 | 450 | 50 |
| 1062 |  |  |  |  |  | 88 | - | 1350 | 1350 | 150 | 0 |
| 1062 |  |  |  |  |  | 93 | 20 | 450 | 150 | 150 | 0 |
| 1062 |  |  |  |  |  | 98 | - | 450 | 450 | 150 | 0 |
| 1062 |  |  |  |  |  | 112 | - | 450 | 450 | 150 | 0 |
| 1062 |  |  |  |  |  | 119 | 0 | 1350 | 150 | 150 | 0 |
| 1062 |  |  |  |  |  | 126 | - | 1350 | 1350 | 50 | 0 |
| 1121 | 28 | F | YES | 4 | YES | 71 | 0 | 1350 | 1350 | 0 | 0 |
| 1121 |  |  |  |  |  | 78 | - | 0 | 0 | 0 | 0 |
| 1121 |  |  |  |  |  | 89 | - | 0 | 0 | 0 | 0 |
| 1121 |  |  |  |  |  | 96 | 0 | 0 | 50 | 0 | 0 |
| 1145 | 39 | F | NO | 2 | YES | 66 | 0 | 1350 | 1350 | 450 | 0 |
| 1145 |  |  |  |  |  | 73 | - | 1350 | 1350 | 450 | 0 |
| 1145 |  |  |  |  |  | 80 | - | 1350 | 450 | 450 | 0 |
| 1145 |  |  |  |  |  | 94 | 40 | 1350 | 1350 | 450 | 50 |
| 1145 |  |  |  |  |  | 101 | - | 1350 | 1350 | 450 | 0 |
| 1145 |  |  |  |  |  | 115 | 20 | 1350 | 1350 | 450 | 0 |
| 1215 | 54 | M | NO | 1 | NO | 79 | 40 | 1350 | 1350 | 450 | 150 |
| 1215 |  |  |  |  |  | 109 | 80 | 1350 | 1350 | 150 | 50 |
| 1234 | 49 | F | NO | 2 | YES | 38 | - |  | 1350 | 4050 | 1350 |
| 1234 |  |  |  |  |  | 44 | - | 1350 | 1350 | 4050 | 150 |
| 1234 |  |  |  |  |  | 52 | - | 1350 | 1350 | 4050 | 450 |
| 1234 |  |  |  |  |  | 60 | - | 1350 | 1350 | 4050 | 150 |
| 1234 |  |  |  |  |  | 68 | - | 1350 | 1350 | 4050 | 150 |
| 1234 |  |  |  |  |  | 82 | - | 1350 | 1350 | 4050 | 150 |
| 1234 |  |  |  |  |  | 94 | - | 1350 | 1350 | 450 | 0 |
| 1278 | 23 | M | NO | 1 | NO | 64 | - | 50 | 1350 | 50 | 0 |
| 1278 |  |  |  |  |  | 74 | - | 1350 | 1350 | 150 | 0 |
| 1278 |  |  |  |  |  | 81 | - | 1350 | 1350 | 150 | 0 |
| 1278 |  |  |  |  |  | 95 | - | 1350 | 450 | 150 | 0 |
| 1278 |  |  |  |  |  | 116 | - | 1350 | 450 | 150 | 0 |
| 1278 |  |  |  |  |  | 130 | - | 1350 | 450 | 150 | 0 |
| 1288 | 27 | F | NO | 2 | YES | 47 | - | 1350 | 1350 | 1350 | 150 |
| 1288 |  |  |  |  |  | 54 | - | 1350 | 1350 | 1350 | 150 |
| 1288 |  |  |  |  |  | 61 | - | 1350 | 1350 | 1350 | 150 |
| 1288 |  |  |  |  |  | 68 | - | 1350 | 450 | 450 | 0 |
| 1288 |  |  |  |  |  | 81 | 80 | 1350 | 1350 | 450 | 50 |
| 1344 | 32 | F | NO | 1 | NO | 63 | 20 | 1350 | 450 | 450 | 50 |
| 1344 |  |  |  |  |  | 105 | 0 | 1350 | 1350 | 150 | 0 |
| 1401 |  |  |  |  |  | 87 | - | 0 | 0 | 1350 | 150 |
| 1401 |  |  |  |  |  | 102 | - | 1350 | 450 | 1350 | 50 |
| 1432 | 54 | M | NO | 1 | NO | 58 | 80 | 1350 | 1350 | 1350 | 150 |
| 1432 |  |  |  |  |  | 84 | 40 | 1350 | 450 | 450 | 50 |
| 1457 | 23 | M | NO | 1 | NO | 80 | - | 1350 | 1350 | 150 | 0 |
| 1457 |  |  |  |  |  | 87 | - | 450 | 150 | 150 | 0 |
| 1457 |  |  |  |  |  | 94 | - | 1350 | 1350 | 450 | 50 |
| 1457 |  |  |  |  |  | 101 | - | 450 | 1350 | 50 | 0 |
| 1457 |  |  |  |  |  | 108 | - | 1350 | 450 | 150 | 0 |
| 1457 |  |  |  |  |  | 122 | - | 1350 | 1350 | 150 | 0 |
| 1462 | 70 | M | NO | 1 | NO | 72 | - | 1350 | 1350 | 450 | 50 |
| 1462 |  |  |  |  |  | 79 | 80 | 1350 | 1350 | 450 | 50 |
| 1462 |  |  |  |  |  | 86 | - | 450 | 450 | 450 | 50 |
| 1462 |  |  |  |  |  | 93 | - | 1350 | 450 | 450 | 50 |
| 1462 |  |  |  |  |  | 100 | 20 | 150 | 1350 | 450 | 0 |
| 1462 |  |  |  |  |  | 107 | - | 1350 | 450 | 150 | 0 |
| 1462 |  |  |  |  |  | 114 | 0 | 450 | 1350 | 450 | 50 |
| 1499 | 42 | M | NO | 1 | NO | 79 | 20 | 1350 | 1350 | 450 | 50 |
| 1499 |  |  |  |  |  | 94 | 0 | 1350 | 450 | 450 | 0 |
| 1551 | 50 | M | YES | 5 | YES | 65 | - | 1350 | 1350 | 4050 | 450 |
| 1551 |  |  |  |  |  | 70 | 320 | 1350 | 1350 | 4050 | 450 |
| 1551 |  |  |  |  |  | 76 | - | 1350 | 1350 | 1350 | 150 |
| 1551 |  |  |  |  |  | 83 | - | 1350 | 1350 | 4050 | 450 |
| 1551 |  |  |  |  |  | 89 | 320 | 1350 | 1350 | 4050 | 450 |
| 1551 |  |  |  |  |  | 97 | - | 1350 | 1350 | 0 | 0 |
| 1551 |  |  |  |  |  | 104 | - | 1350 | 1350 | 1350 | 150 |
| 1551 |  |  |  |  |  | 111 | 160 | 1350 | 1350 | 450 | 150 |
| 1678 | 48 | F | NO | 2 | YES | 54 | - | 1350 | 1350 | 450 | 150 |
| 1678 |  |  |  |  |  | 85 | - | 1350 | 1350 | 1350 | 450 |
| 1678 |  |  |  |  |  | 92 | - | 1350 | 1350 | 1350 | 150 |
| 1817 | 28 | F | NO | 1 | NO | 44 | - | 1350 | 1350 | 450 | 150 |
| 1817 |  |  |  |  |  | 51 | - | 1350 | 1350 | 1350 | 450 |
| 1817 |  |  |  |  |  | 63 | - | 1350 | 1350 | 1350 | 450 |

DPO Days post onset of symptoms; S/ECD Spike ectodomain; S/RBD Spike receptor-binding domain; VN Virus neutralization

**Supplementary Table 2:** Predictive values and likelihood ratios of the ELISA methods as a rogate for virus neutralizing antibody titer of ≥160.

| DPO | Effect | S/ECD $\geq 1350$ | | S/RBD $\geq 1350$ | | S/RBD IgG $\geq 1350$ | | S/RBD IgM $\geq 450$ | |
| --- | --- | --- | --- | --- | --- | --- | --- | --- | --- |
|  |  | Value | 95% CI | Value | 95% CI | Value | 95% CI | Value | 95% CI |
| Overall<br>0-142 | PPV | 0.52 | 0.46 to 0.59 | 0.57 | 0.50 to 0.63 | 0.72 | 0.65 to 0.79 | 0.38 | 0.30 to 0.46 |
|  | NPV | 0.75 | 0.65 to 0.83 | 0.87 | 0.78 to 0.92 | 0.82 | 0.75 to 0.87 | 0.90 | 0.84 to 0.94 |
|  | LR+ | 1.34 |  | 1.61 |  | 3.18 |  | 3.72 |  |
|  | LR- | 0.40 |  | 0.19 |  | 0.26 |  | 0.69 |  |
| 1-30 | PPV | 0.92 | 0.65 to 1.00 | 0.94 | 0.72 to 0.99 | 0.78 | 0.55 to 0.91 | 1.00 | 0.68 to 1.00 |
|  | NPV | 0.67 | 0.47 to 0.82 | 0.80 | 0.58 to 0.92 | 0.65 | 0.43 to 0.82 | 0.56 | 0.39 to 0.73 |
|  | LR+ | 9.85 |  | 13.43 |  | 2.83 |  | - |  |
|  | LR- | 0.45 |  | 0.22 |  | 0.44 |  | 0.62 |  |
| 31-60 | PPV | 0.79 | 0.67 to 0.88 | 0.69 | 0.57 to 0.78 | 0.73 | 0.61 to 0.82 | 0.90 | 0.74 to 0.96 |
|  | NPV | 0.56 | 0.41 to 0.70 | 0.72 | 0.52 to 0.83 | 0.72 | 0.55 to 0.84 | 0.55 | 0.43 to 0.66 |
|  | LR+ | 1.80 |  | 1.56 |  | 1.92 |  | 6.23 |  |
|  | LR- | 0.38 |  | 0.28 |  | 0.28 |  | 0.59 |  |
| 61-90 | PPV | 0.32 | 0.23 to 0.43 | 0.35 | 0.25 to 0.47 | 0.58 | 0.41 to 0.72 | 0.50 | 0.31 to 0.69 |
|  | NPV | 1.00 | 0.65 to 1.00 | 1.00 | 0.74 to 1.00 | 0.90 | 0.79 to 0.96 | 0.78 | 0.66 to 0.86 |
|  | LR+ | 1.13 |  | 1.30 |  | 3.20 |  | 2.40 |  |
|  | LR- | 0.00 |  | 0.00 |  | 0.27 |  | 0.69 |  |
| 91-120 | PPV | 0.52 | 0.40 to 0.64 | 0.61 | 0.48 to 0.73 | 0.87 | 0.71 to 0.95 | 0.80 | 0.49 to 0.96 |
|  | NPV | 1.00 | 0.77 to 1.00 | 1.00 | 0.85 to 1.00 | 0.87 | 0.74 to 0.94 | 0.62 | 0.50 to 0.73 |
|  | LR+ | 1.43 |  | 2.05 |  | 8.80 |  | 5.21 |  |
|  | LR- | 0.00 |  | 0.00 |  | 0.20 |  | 0.79 |  |
| >120 | PPV | 0.43 | 0.16 to 0.75 | 0.43 | 0.16 to 0.75 | 0.75 | 0.30 to 0.99 | 0.00 | 0.00 to 0.95 |
|  | NPV | 1.00 | 0.05 to 1.00 | 1.00 | 0.05 to 1.00 | 1.00 | 0.51 to 1.00 | 0.57 | 0.25 to 0.84 |
|  | LR+ | 1.25 |  | 1.25 |  | 5.00 |  | 0.00 |  |
|  | LR- | 0.00 |  | 0.00 |  | 0.00 |  | 1.25 |  |

DPO Days post onset of symptoms; S/ECD Spike ectodomain; S/RBD Spike receptor-binding domain; PPV Positive predictive value; NPV Negative predictive value; LR+ Positive likelihood ratio; LR- Negative likelihood ratio

## References

1. Salazar, E., et al. Relationship between Anti-Spike Protein Antibody Titers and SARS-CoV-2 In Vitro Virus Neutralization in Convalescent Plasma. bioRxiv (2020).

2. Iyer, A.S., et al. Dynamics and significance of the antibody response to SARS-CoV-2 infection. medRxiv, 2020.2007.2018.20155374 (2020).

3. Wajnberg, A., et al. SARS-CoV-2 infection induces robust, neutralizing antibody responses that are stable for at least three months. medRxiv, 2020.2007.2014.20151126 (2020).

4. Ibarrondo, F.J., et al. Rapid Decay of Anti–SARS-CoV-2 Antibodies in Persons with Mild Covid-19. New England Journal of Medicine (2020).

5. Long, Q.-X., et al. Clinical and immunological assessment of asymptomatic SARS-CoV-2 infections. Nature Medicine 26, 1200–1204 (2020).

6. Seow, J., et al. Longitudinal evaluation and decline of antibody responses in SARS-CoV-2 infection. medRxiv, 2020.2007.2009.20148429 (2020).

7. Sethuraman, N., Jeremiah, S.S. & Ryo, A. Interpreting Diagnostic Tests for SARS-CoV-2. Jama (2020).

8. Liu, X., et al. Patterns of IgG and IgM antibody response in COVID-19 patients. Emerg Microbes Infect 9, 1269–1274 (2020).

9. Zhu, F.C., et al. Safety, tolerability, and immunogenicity of a recombinant adenovirus type-5 vectored COVID-19 vaccine: a dose-escalation, open-label, non-randomised, first-in-human trial. Lancet 395, 1845–1854 (2020).

10. Jackson, L.A., et al. An mRNA Vaccine against SARS-CoV-2-Preliminary Report. N Engl J Med (2020).

11. Bar-Zeev, N. & Moss, W.J. Encouraging results from phase 1/2 COVID-19 vaccine trials. Lancet 396, 448–449 (2020).

12. Folegatti, P.M., et al. Safety and immunogenicity of the ChAdOx1 nCoV-19 vaccine against SARS-CoV-2: a preliminary report of a phase 1/2, single-blind, randomised controlled trial. Lancet 396, 467–478 (2020).

13. Zhu, F.C., et al. Immunogenicity and safety of a recombinant adenovirus type-5-vectored COVID-19 vaccine in healthy adults aged 18 years or older: a randomised, double-blind, placebo-controlled, phase 2 trial. Lancet 396, 479–488 (2020).

14. Tan, C.W., et al. A SARS-CoV-2 surrogate virus neutralization test based on antibody-mediated blockage of ACE2-spike protein-protein interaction. Nat Biotechnol (2020).

15. Salazar, E., et al. Treatment of COVID-19 Patients with Convalescent Plasma Reveals a Signal of Significantly Decreased Mortality. Am J Pathol (2020).

16. Joyner, M.J., et al. Effect of Convalescent Plasma on Mortality among Hospitalized Patients with COVID-19: Initial Three-Month Experience. medRxiv, 2020.2008.2012.20169359 (2020).

17. Lee, N., et al. Anti-SARS-CoV IgG response in relation to disease severity of severe acute respiratory syndrome. J Clin Virol 35, 179–184 (2006).

18. Lynch, K.L., et al. Magnitude and kinetics of anti-SARS-CoV-2 antibody responses and their relationship to disease severity. Clin Infect Dis (2020).

19. Stadlbauer, D., et al. SARS-CoV-2 Seroconversion in Humans: A Detailed Protocol for a Serological Assay, Antigen Production, and Test Setup. Curr Protoc Microbiol 57, e100 (2020).

20. Okuno, T. & Kondelis, N. Evaluation of dithiothreitol (DTT) for inactivation of IgM antibodies. J Clin Pathol 31, 1152–1155 (1978).

21. Sui, J., et al. Potent neutralization of severe acute respiratory syndrome (SARS) coronavirus by a human mAb to S1 protein that blocks receptor association. Proc Natl Acad Sci U S A 101, 2536–2541 (2004).

22. R Development Core Team. R: a language and environment for statistical computing. (R Foundation for Statistical Computing, Vienna, Austria, 2014).

23. Therneau, T.M. A Package for Survival Analysis in R (Springer, New York, 2020).

24. Kassambara and Kosinski. “Survminer: Drawing Survival Curves using “ggplot2”. (2018).

